# Application of machine learning in the discovery of antimicrobial peptides: Exploring their potential for ulcerative colitis therapy

**DOI:** 10.1101/2025.06.17.660148

**Authors:** Hui Miao, Ziwei Wang, Shihu Chen, Jiaqi Wang, Hongyue Ma, Yifan Liu, Hui Yang, Ziyi Guo, Jiamei Wang, Pengfei Cui

**Affiliations:** Lab of Environmental Health and Ecological Engineering, College of Marine Life Science, Ocean University of China, Qingdao 266003, China; Haide College, Ocean University of China, Qingdao, China

**Keywords:** Antimicrobial peptide, antimicrobial peptide prediction, ulcerative colitis, gut microbiota, intestinal barrier damage, *Akkermansia muciniphila*

## Abstract

Ulcerative colitis (UC) is a chronic inflammatory bowel disease with rising global prevalence, yet existing treatments are not universally effective. Antimicrobial peptides (AMPs), produced by the immune system, have diverse antimicrobial and immune-regulatory functions, making them promising candidates for UC therapy. Using machine learning, we developed a machine learning-based prediction model to identify novel AMPs. The predicted peptides demonstrated significant biological activity in vitro and in vivo. In a dextran sulfate sodium-induced UC mouse model, engineered AMPs notably improved UC-related parameters, such as body weight, disease activity index (DAI), and colon length. These effects were likely mediated by modulation of *Akkermansia muciniphila*. This study highlights the potential of machine learning-identified AMPs as future therapeutic candidates for UC.

**Graphical abstract:** 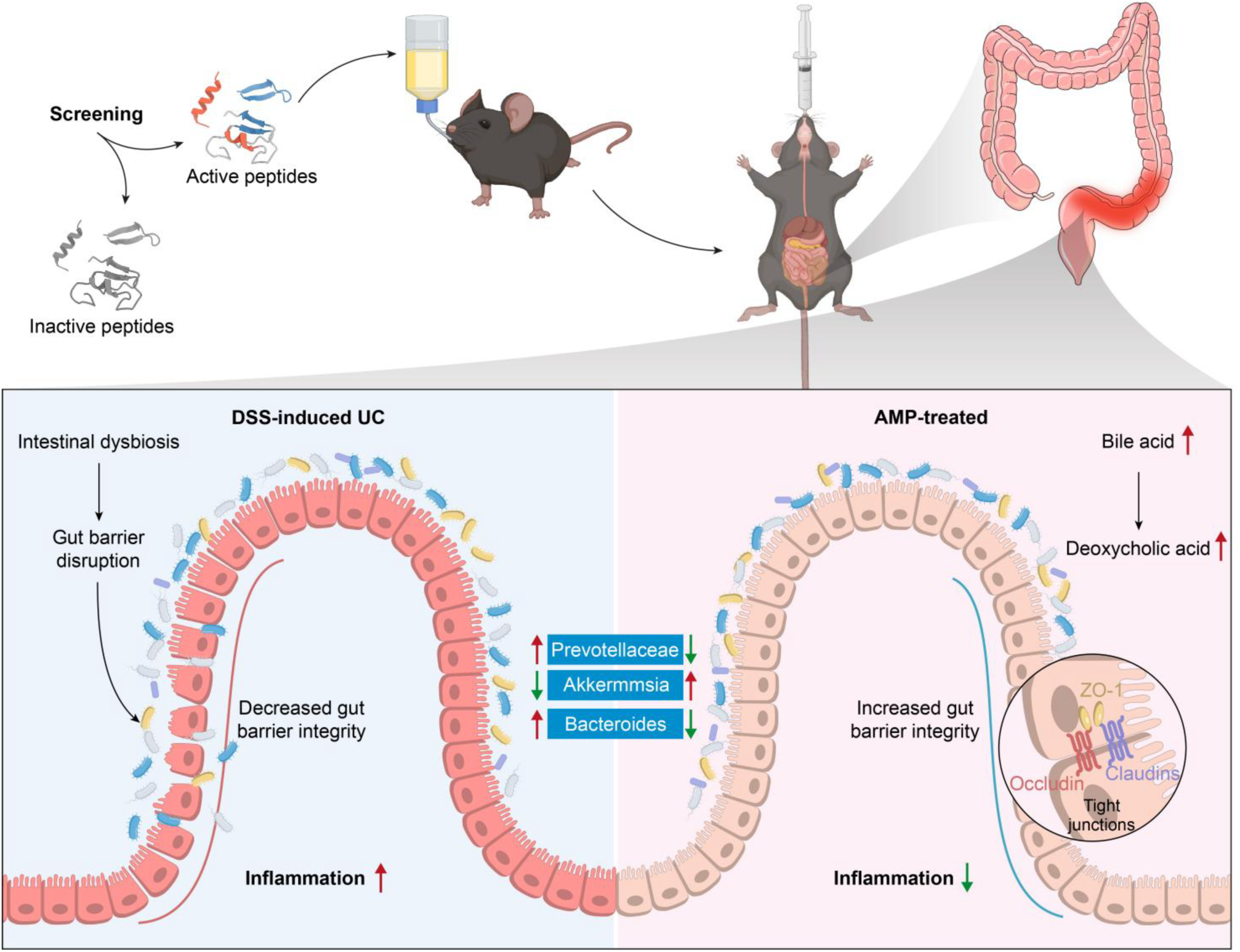

## Introduction

Ulcerative colitis (UC) is a chronic, recurrent inflammatory bowel disease (IBD) that predominantly affects the colon and rectum.^1–2^ Its precise pathogenesis remains elusive, while its rapidly increasing global prevalence has become a significant healthcare challenge.^3^ UC is characterized by symptoms such as abdominal pain, diarrhea, hematochezia, etc.^4^ Moreover, its complexity often leads to severe complications, including intestinal bleeding, toxic megacolon, perforation, and a heightened risk of colorectal cancer. In addition, environmental factors, genetic susceptibility, dysregulation of immune response and dysregulation of gut microbiome are considered to be the most important factors in the formation of UC.^5–6^ Despite advances in treatment techniques, including 5-aminosalicylic acid,^7^ antibiotics, immunosuppressants, and biologics (such as infliximab and Adalimumab) have been used clinically, there remains no universally effective cure and no specific drugs for the treatment of this disease.^8–9^ Additionally, these treatments are not universally applicable, and long-term use of these drugs can lead to a range of side effects,^10^ including liver and kidney toxicity, risk of drug resistance, autoimmune reactions, viral infections, and tumorigenesis.^11–13^ Therefore, there is an urgent need to seek novel therapeutic strategies for UC treatment. The gut microbiome constitutes a complex ecosystem that plays a pivotal role in the host’s digestion, nutrient absorption, and immune regulation.^14^ Dysbiosis, or imbalances in the composition and function of the gut microbiota, can disrupt intestinal homeostasis, leading to various diseases.^15^ Therefore, intestinal microbiome imbalance may be one of the reasons for UC induction.

The immune system contributes to maintaining a balanced host-microbe relationship, primarily through the production of immunoglobulin A (IgA) and the induction of antimicrobial peptides (AMPs).^16^ AMPs, also known as host defense peptides (HDPs),^17^ are small, naturally occurring amphipathic molecules, typically comprising 12 to 50 amino acid residues.^18^ These peptides exhibit a net positive charge under physiological conditions, which enables them to interact with microbial membranes.^18^ AMPs possess broad-spectrum antimicrobial activity, along with additional properties such as promoting wound healing and adsorbing lipopolysaccharides (LPS).^18^

As key effectors of the innate immune system and the first line of defense against pathogenic infections, AMPs hold significant clinical potential.^19–22^ However, their application remains limited, particularly in the treatment of conditions like UC where few studies have yet explored the use of AMPs as a therapeutic approach. This limited use is partly due to the challenges in reliably identifying AMPs and balancing their antimicrobial efficacy with cytotoxicity. Current methods for discovering AMPs primarily rely on extraction from living organisms, a process that is both time-consuming and unpredictable, often necessitating repeated experimental validation.^23–25^ To efficiently and cost-effectively identify effective antimicrobial peptides (AMPs), we employed computational and bioinformatics approaches to explore the relationship between the biological properties and antimicrobial activity of AMPs, leveraging extensive datasets for inferences. This method circumvents the need for biological experiments, relying instead on robust algorithms and the high computational power of modern systems to perform predictive analyses. Consequently, we constructed a machine learning model trained on multiple datasets to screen for safe and potent AMPs. In this study, we also examined the therapeutic effects of the identified AMPs in ulcerative colitis (UC), comparing their efficacy with that of the standard treatment, 5-aminosalicylic acid, and the antibiotic ciprofloxacin in glucan-induced UC mouse models. To further investigate the mechanisms, we used 16S rRNA sequencing to analyze the impact of AMPs on the intestinal microbiota and employed metabolomics to assess changes in metabolite levels in fecal samples, providing insights into the potential mechanisms by which AMPs exert their effects. We hypothesized that AMPs could treat UC by modulating the inflammatory microenvironment, clearing lipopolysaccharides (LPS), and restoring gut microbiota balance. Our in vitro and in vivo validation experiments support this hypothesis, showing that the designed AMPs can effectively treat UC through anti-inflammatory actions and microbiome modulation.

## Results

### Establishment of machine learning model trained by data set

During the hyperparameter optimization process, we meticulously adjusted two key models: the Antimicrobial Peptide Predictor (Peptide_predictor) and the Toxicity Predictor (Toxic_predictor). The performance of the optimized models was significantly enhanced, as evidenced by the results presented in Figure 1A and Figure 1B.

**Figure 1.**
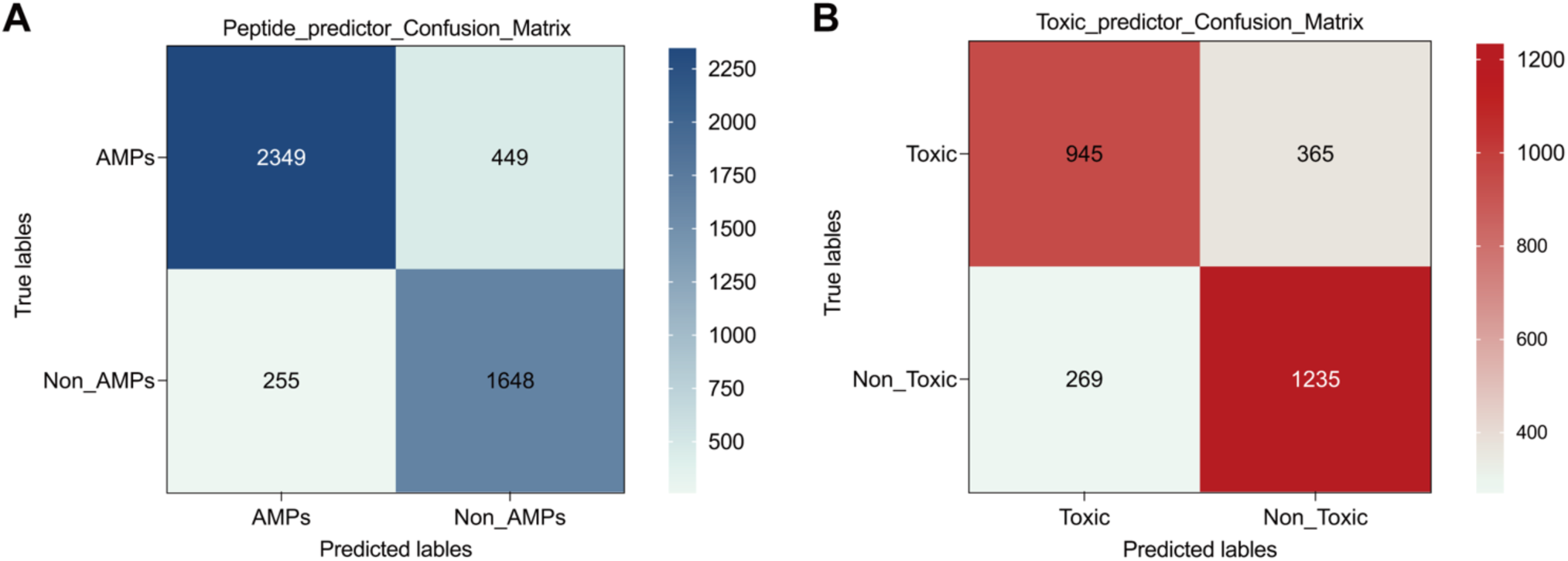
Establishment of machine learning model based on data set training. (A) Confusion Matrix of the Peptide_predictor Model. (B) Confusion Matrix of the Toxic_predictor Model.

Post-optimization, the Peptide_predictor model exhibited superior performance in identifying AMPs, with its F1 score improving to 0.85. These substantial improvements in the metrics underscore the increased reliability of the model in predicting the presence or absence of AMPs.

For the Toxic_predictor model, despite certain deficiencies in toxicity prediction, the optimized F1 score was improved to 0.75. Analysis of the confusion matrix revealed a bias in the model’s ability to distinguish true positives from false negatives. This bias may be attributable to a correlation between the toxicity of AMPs and their bactericidal efficacy, which could partly explain the model’s suboptimal performance in toxicity prediction.

Further investigation supports the existence of a correlation between the toxicity of AMPs and their bactericidal capacity. Through a comprehensive review of the literature,^40^ we have observed a notable consistency in the distribution of toxicity and bactericidal activity among AMPs. This observation elucidates the intrinsic relationship between toxicity and bactericidal efficacy, while also providing an explanation for the model’s suboptimal performance.

Through the application of a genetic algorithm, we evaluated 6,373 AMPs sequences over 10 iterative rounds, ultimately identifying 22 target sequences based on our predefined selection criteria (refer to the attached table). This methodology not only enhanced screening efficiency but also augmented the likelihood of discovering novel AMPs.

### Modified synthesis and characterization of AMPs

To increase the positive charge, serine residues were substituted with positively charged lysine and arginine Additionally, tryptophan and proline were incorporated to replace certain amino acids in the original AMPs, thereby enhancing hydrophobicity. Amidation at the C-terminal was employed to improve selectivity between different bacterial cells. The following five AMPs were synthesized, as detailed in Table 1. The molecular weights of these AMPs were confirmed via mass spectrometry (MS). Table 2 and Figure S1 present the relevant properties of the modified AMPs. The experimentally determined molecular weights closely matched the theoretical values, and the purity exceeded 95%, confirming the successful synthesis of LR, KC, KI, KP, and WR. The net charges of the AMPs ranged from +6 to +7, while their hydrophobicity varied between 40% and 61%. Additionally, the secondary structures of these AMPs were predicted using AlphaFold3. The results indicated that LR, KC, KP, and WR exhibited α-helix structures, whereas KI demonstrated random coiling (Figure 2).

**Figure 2.**
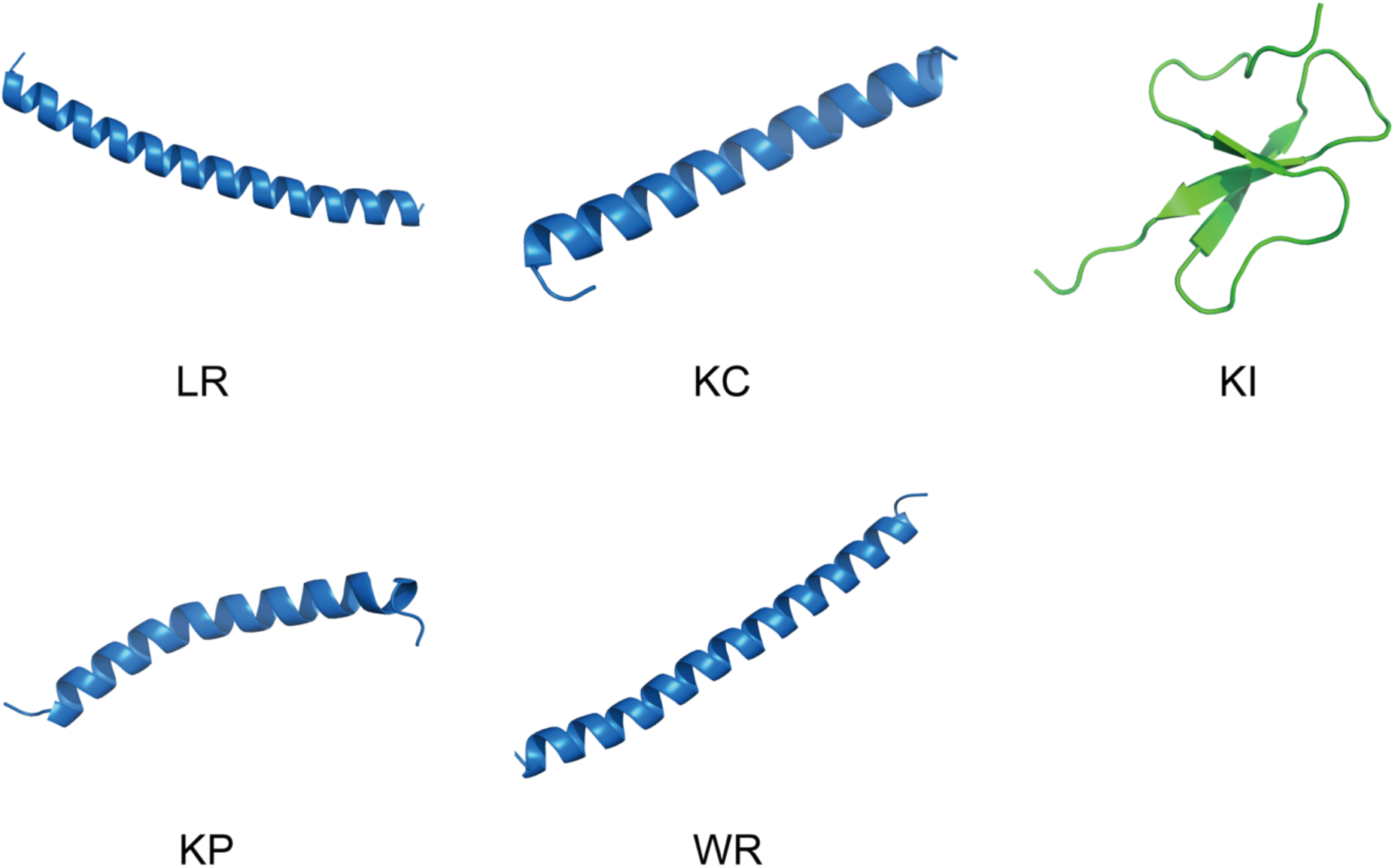
Alphafold3 predicts the secondary structure outcomes of these five AMPs.

**Table 1.**
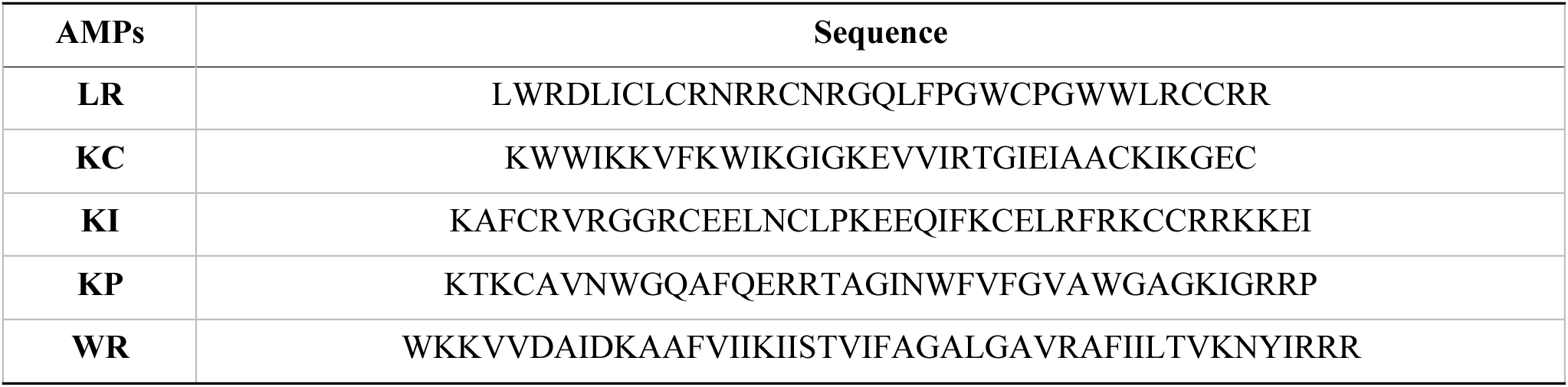
Sequence of AMPs.

**Table 2.** The relevant attribute parameters of the designed AMPs.

| AMPs | M. Wt. (actual value) | M. Wt. (theoretical value) | net charge | hydrophobic ratio (%) | purity (%) | Probability_score | sterilize_proba |
| --- | --- | --- | --- | --- | --- | --- | --- |
| LR | 4293.13 g/mol | 4294.11 g/mol | 7 | 50% | 95.12% | 0.99997103 | 0.463136199 |
| KC | 4058.98 g/mol | 4059.95 g/mol | 6 | 51% | 95.38% | 0.566047 | 0.626558879 |
| KI | 4903.87 g/mol | 4904.84 g/mol | 7 | 40% | 95.86% | 0.99985147 | 0.630466238 |
| KP | 4263.89 g/mol | 4264.86 g/mol | 6 | 45% | 95.24% | 0.99964035 | 0.635288829 |
| WR | 5158.27 g/mol | 5159.23 g/mol | 7 | 61% | 95.43% | 0.9999997 | 0.373250428 |

Furthermore, we employed this algorithm to predict the accuracy and toxicity of modified AMPs. The fingdings revealed that the accuracy for LR, KC, KI, KP, and WR exceeded 50%. Specifically, the toxicity prediction results were 46.31% for LR, 62.65% for KC, 63.04% for KI, and 63.5% for KP, with WR having a 37.35% probability of being toxic (Table 2).

### In vitro antibacterial activity and biocompatibility of AMPs

To validate the algorith200 μg/ml did not exhibit cytotoxic effects on RAW264.7 cells. In contrast, the viability of RAW264.7 cells progressively declined with increasing concentrations of Km’s accuracy, we conducted in vitro bactericidal and biocompatibility assays of AMPs. The findings indicated that LR and KC exhibited the most effective bactericidal activity among the five AMPs tested, achieving a 100% bactericidal rate at a concentration of 25 μg/ml. Subsequently, we evaluated the biocompatibility of the five AMPs: LR, KC, KI, KP, and WR. The cytotoxicity assay using CCK-8 indicated that LR concentrations ranging from 0 to C within the same range, with cell viability reduced to only 20.44% at 200 μg/ml. This suggests that KC exhibits significantly higher cytotoxicity compared to LR. Furthermore, the cytotoxic effects of KI, KP, and WR were observed to increase with rising concentrations, displaying greater cytotoxicity than LR but lower than KC (Figure 3B). Upon analyzing the outcomes of bactericidal and toxicity assays, we opted to select LR and KC, which demonstrated superior results, for subsequent experimentation.

**Figure 3.**
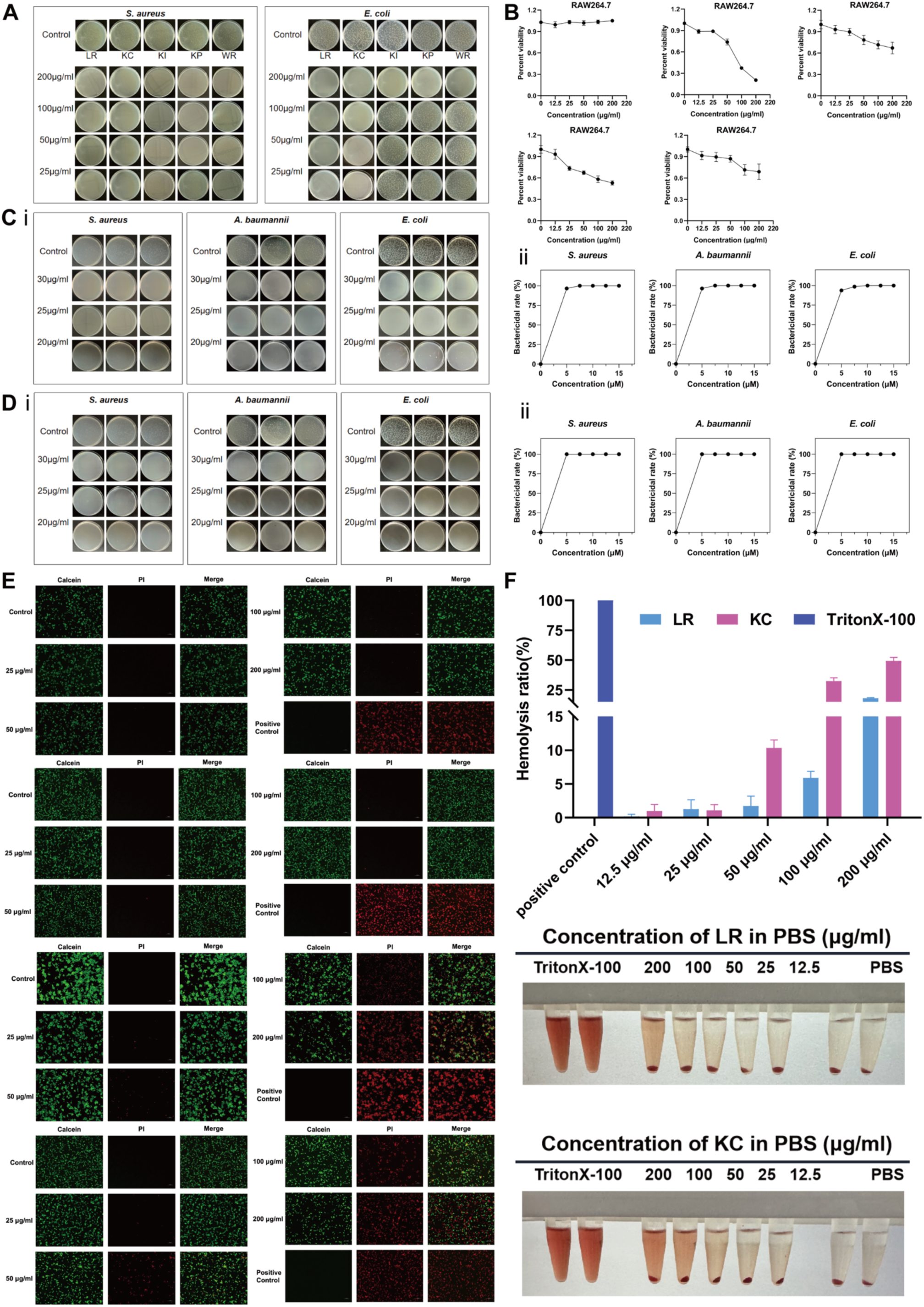
In vitro experiment of AMPs. (A) Bactericidal effect of five AMPs on two kinds of bacteria. (B) The cytotoxicity of five AMPs to macrophages was LR, KC, KI, KP,WR, respectively. (C) The bactericidal effect of LR against *S. aureus*, *AB-29* and *E. coli*. (D) The bactericidal effect of KC against *S. aureus*, *AB-29* and *E. coli*. (E) Cytotoxicity of LR and KC to macrophages and colon cancer cells. (F) The hemolytic activity of LR and KC. Data are presented as mean ± SD (n = 3).

Consequently, we chose *E. coli*, *S. aureus*, and resistant Acinetobacter baumannii to assess the Minimum Bactericidal Concentration (MBC) of AMPs. The findings indicated that the bactericidal rates of LR against *E. coli*, *S. aureus* and resistant *A. baumannii* were 96.6%, 96.3% (Figure 3C), and 93.8% at a concentration of 20 μg/ml, respectively. In contrast, the bactericidal rates of KC against these bacterial strains were 100% at the same concentration (Figure 3D).

Simultaneously, the cell staining results indicated that the cytotoxicity of KC was significantly higher than that of LR, with an increase in the number of dead cells correlating with higher KC concentrations (Figure 3E). The assessment of the hemolytic activity of AMPs revealed that, within the MBC range, the hemolytic rates for LR and KC were below 3%. At a concentration of 100 μg/ml, LR exhibited a hemolytic rate of less than 8%, indicating slight hemolysis, whereas KC demonstrated a hemolytic rate of less than 34%, indicating substantial hemolysis (Figure 3F).

### In vitro evaluation of anti-inflammatory effect of AMPs and results of CD spectrum

Initially, lipopolysaccharide (LPS) at concentrations of 0.1 μg/ml,1 μg/ml, 10 μg/ml and 100 μg/ml was employed to stimulate RAW264.7 cells. Subsequently, cell viability was seeessed using the CCK-8 assay. The findings indicated a cell survival rate of 91.4% at 0.1 μg/ml, with a significant decline in survival rate observed as LPS concentration increased (Figure 4A). An inflammation model was established using LPS at 0.1 μg/ml. Following this, varying concentration gradients of LR and KC were introduced, and the levels of inflammatory cytokines IL-6 and TNF-α were quantified using an ELISA kit (Jiang-lai Bio). The results indicated a decrease in the levels of inflammatory factors with increasing concentrations of LR, as demonstrated in Figure 4B. Specifically, LR at a concentration of 200 μg/ml significantly reduced the levels of inflammatory factors compared to the LPS group. In contrast, KC at the same concentration did not exhibit a significant effect on the reduction of inflammatory factors (Figure 4C). Consequently, it can be inferred that LR possesses a certain degree of anti-inflammatory activity relative to KC. Considering the aforementioned bactericidal, cytotoxic, hemolytic, and anti-inflammatory findings, LR was ultimately selected for subsequent experiments.

**Figure 4.**
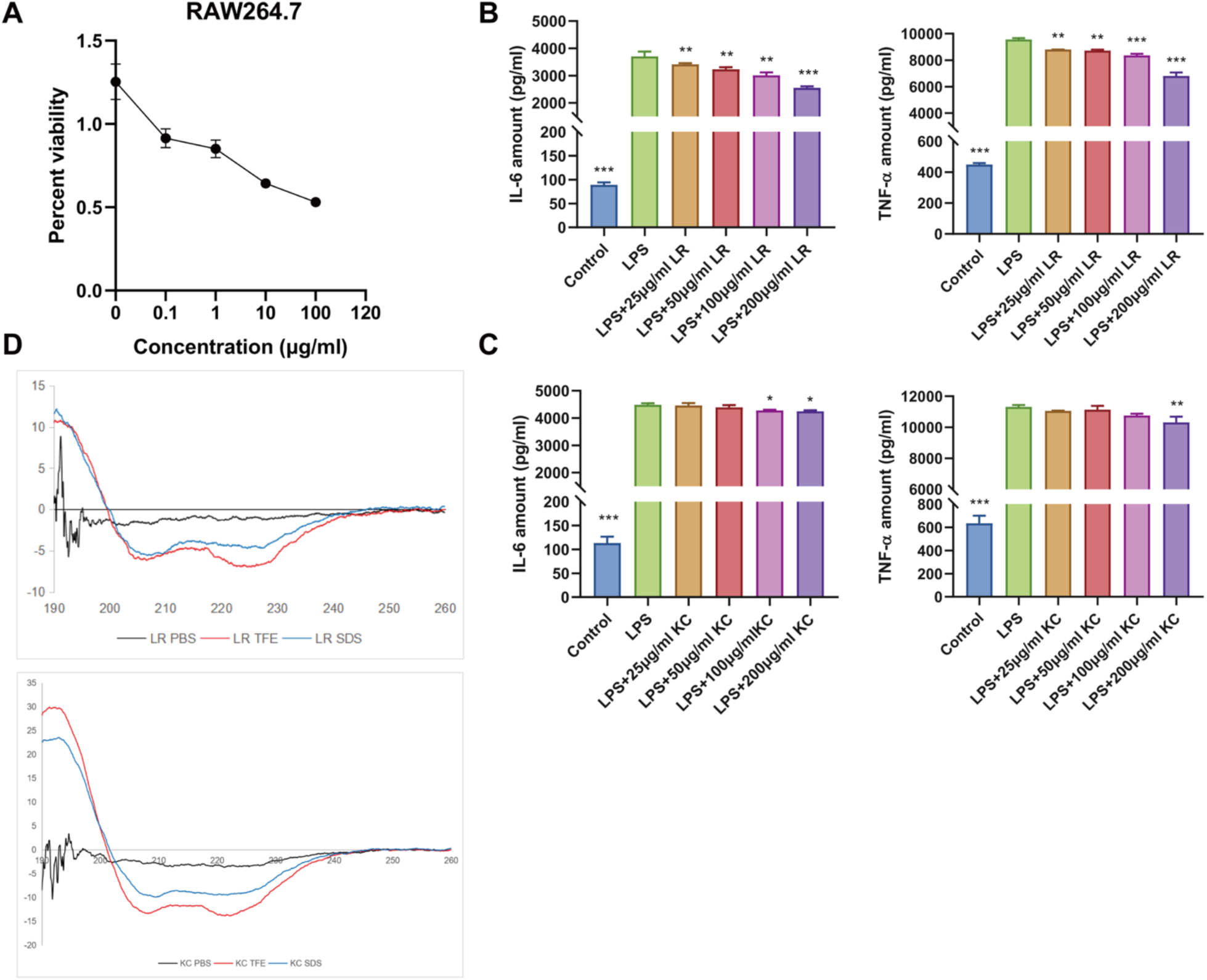
Anti-inflammatory effects of LR and KC and their CD experiments. (A) Cytotoxicity of LPS to macrophages (B) Effects of LR on levels of inflammatory cytokines IL-6 and TNF-α. (C) Effects of KC on levels of inflammatory cytokines IL-6 and TNF-α. (D) CD spectral results of LR and KC in PBS, SDS and TFE solutions. Data are presented as mean ± SD (n = 3); * means p < 0.05, ** means p < 0.01, *** means p < 0.001.

Furthermore, the circular dichroism (CD) spectroscopy data (Figure 4D) indicate that the LR and KC spectra in 10 mM PBS at pH7.4 exhibited unordered conformations, lacking the formation of α-helix structures. In contrast, in a 50% TFE solution and a 30 mM anionic SDS solution, both LR and KC demonstrated α-helical structures, as evidenced by signals at 208 nm and 220 nm.

### In vitro stability of AMPs

The results from the *in vitro* thermal stability experiments revealed that, at 60℃, the bactericidal rate remained at 100% when the concentration of AMPs was 30 μg/ml, 40 μg/ml, and 50 μg/ml. The findings suggests that the bactericidal activity of AMPs was consistent before and after thermal treatment (Figure 5A).

**Figure 5.**
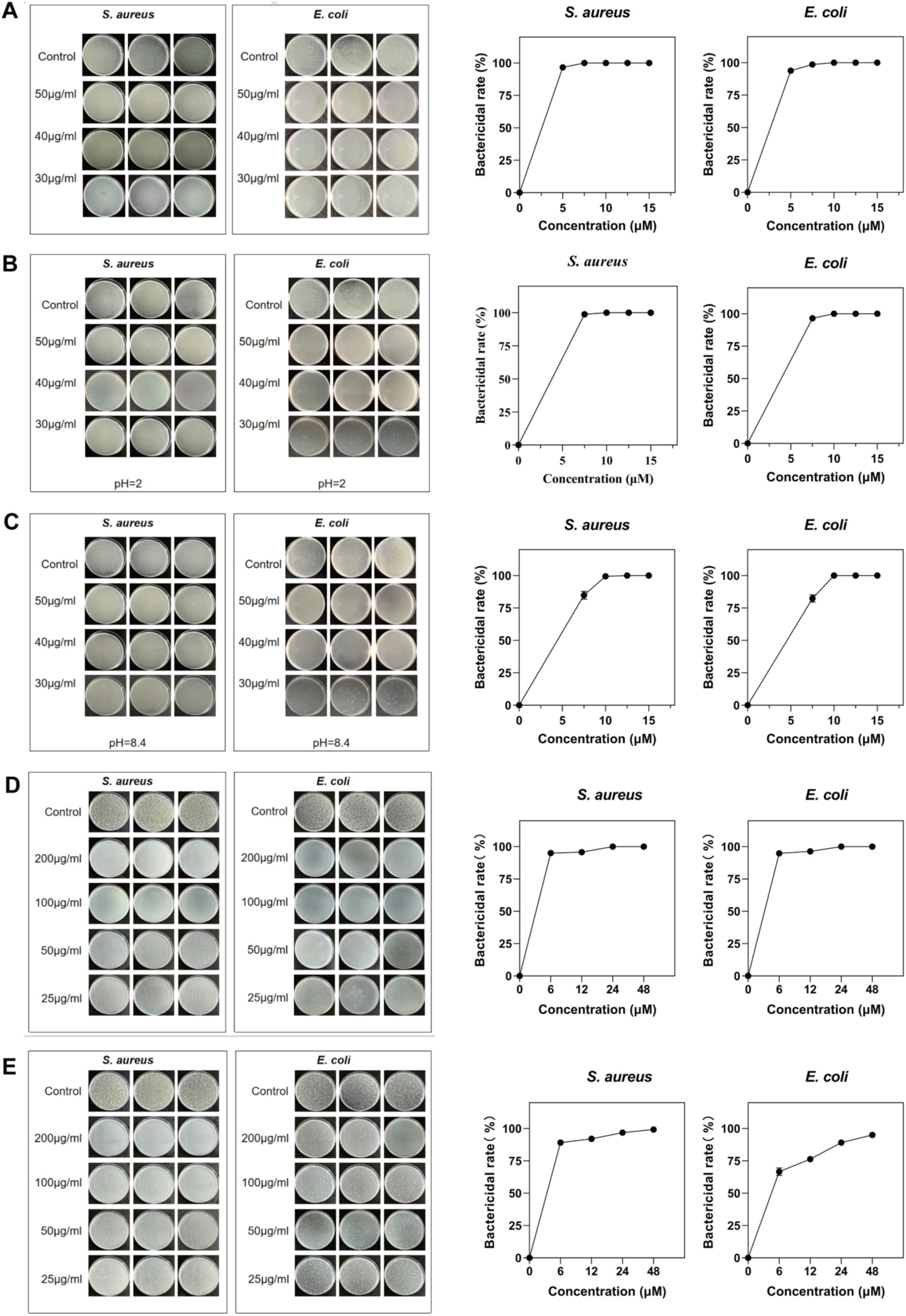
Stability of simulated AMPs in vivo. (A) LR bactericidal effect at 60℃. (B) The bactericidal effect of LR after treatment with pH=2 solution. (C) The bactericidal effect of LR after treatment with pH=8.4 solution. (D) The bactericidal effect of LR after pepsin treatment. (E) The bactericidal effect of LR after trypsin treatment.

The in vitro acid-base stability test results demonstrated that following treatment with AMPs in simulated gastric fluid at pH2 (Figure 5B) and simulated intestinal fluid at pH8.4 (Figure 5C), the bactericidal efficacy of LR against Staphylococcus aureus remained consistent before and after exposure to pH2. However, for *E. coli*, the bactericidal rate decreased to 96.5% at a concentration of 30μg/ml. Upon treatment at pH8.4, the bactericidal rates for *E. coli* and *S. aureus* declined from 100% to 82.3% and 84.8%, respectively, at a concentration of 30 μg/ml. These findings suggest that alkaline conditions may adversely affect the efficacy of AMPs at lower concentrations.

The in vitro proteolytic stability assessment of AMPs revealed that following pepsin treatment, the bactericidal efficacy of LR remained at 100% at a concentration of 100 μg/ml. However, at a reduced concentration of 50 μg/ml, the bactericidal rate of LR decreased to approximately 95% for both *E. coli* and *S. aureus*, as illustrated in Figure 5D. Furthermore, post-trypsin treatment (Figure 5E) resulted in a decline in the bactericidal rate against *S. aureus* from 100% to 92% at 50 μg/ml, while for *E. coli*, it decreased to 75%. These findings suggest that the bactericidal activity of AMPs is somewhat compromised following trypsin treatment.

### Effects of DSS and various therapeutic drugs on clinical symptoms

Mice were administered DSS water ad libitum for a duration of seven days to induce a model of ulcerative colitis (UC). The spleen index and colon length serve as critical indicators of the UC disease phenotype. Treatment with DSS water led to significant weight loss (Figure 6A), an increase DAI (refer to Figure 6B), a reduction in colon length, and an elevated spleen index in the mice. Based on these observations, it was concluded that the UC model was successfully established. Subsequently, experimental groups receiving AMPs, 5-ASA, and CIP groups were designed to evaluate the effects of various therapeutic agents on colitis.Among these, the AMPs group demonstrated the most pronounced therapeutic efficacy when compared to the CIP and 5-ASA groups. In comparison to the DSS group, the body weight of mice progressively increased starting from the second day of gavage treatment, while both the DAI score and spleen index gradually decreased throughout the treatment period (Figure 6C-E). The administration of AMPs via gavage demonstrated the most pronounced effects, resulting in the absence of occult blood in the stool and body weight closely resembling that of the healthy. control group by the final day of treatment. In comparison to the control group, which exhibited a spleen index of 2.22±0.23 mg/g, the DSS group demonstrated a significantly elevated spleen index of 5.85±1 mg/g. Conversely, the spleen indices for the AMPs group (3.67±0.5 mg/g), the 5-ASA group (4.29±0.25 mg/g) and the CIP group (5.22±0.5 mg/g) were comparatively lower.

**Figure 6.**
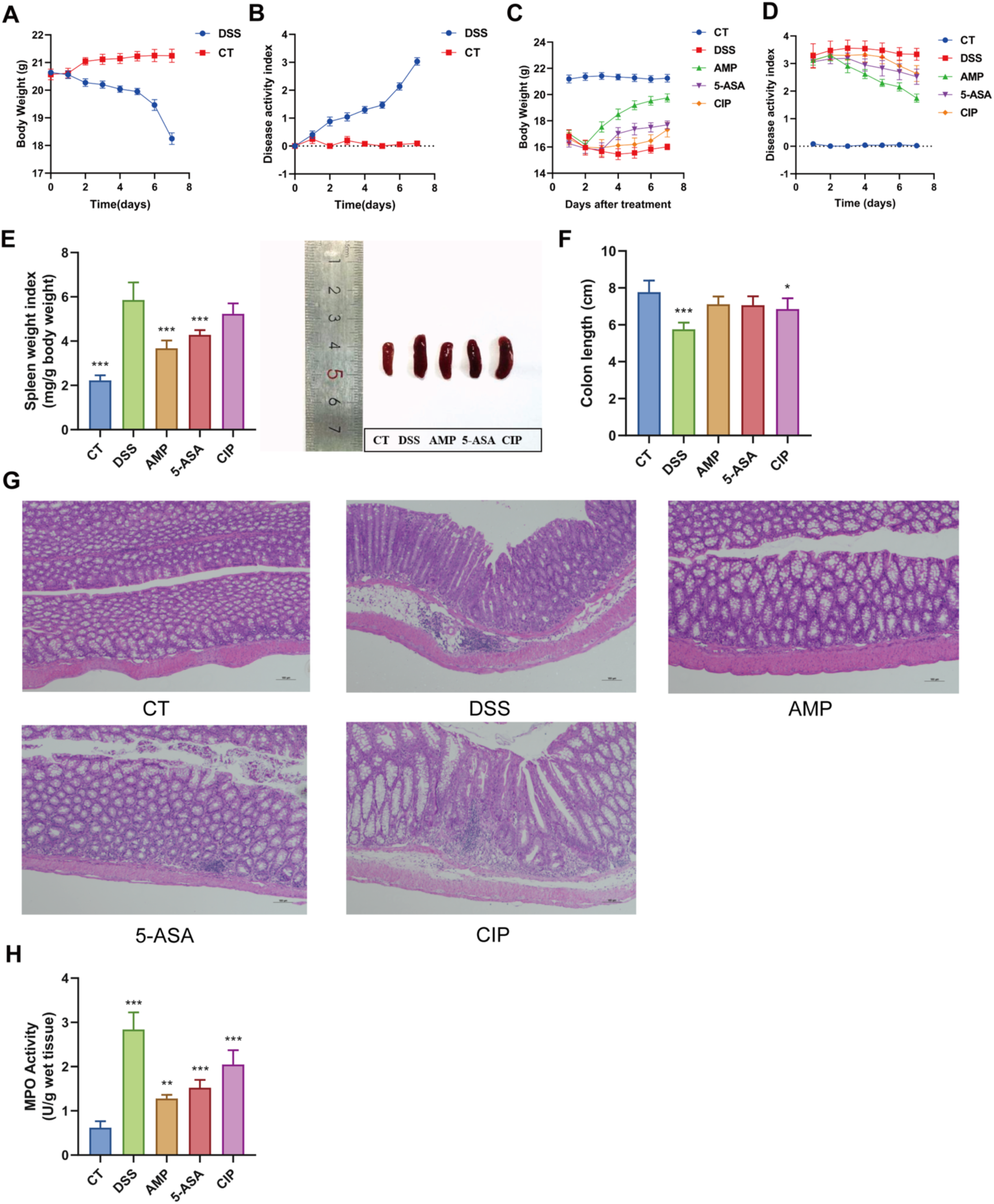
The influence of various therapeutic drugs on clinical symptoms. (A) Changes of body weight in mice after DSS. (B) The change of DAI score after DSS Data are means ± SEM, n = 6 in DSS group and in the other group. (C) Changes in body weight of mice after drug treatment. (D) Changes of DAI score in each group of mice after treatment. (E) Changes of spleen index in mice after drug treatment. Significant differences were indicated by *p < 0.05, **p < 0.01, ***p < 0.001 (F) length of colon between the ileocecal junction and the proximal rectum. (G) Representative images of the colon obtained by H&E staining (100x) were obtained in the CT, DSS, AMPs, 5-ASA, and CIP groups. (H) MPO content in colon tissues of each treatment group. Data are presented as mean ± SD (n = 6); * means p < 0.05, ** means p < 0.01, *** means p < 0.001.

Similarly, the colons of the DSS group were notably shorter, measuring 5.75±0.55 cm, compared to the control group, which measured 7.78±0.5 cm. However, post-treatment recovery was observed in the colon lengths of the AMPs group (7.12±0.82 cm), the 5-ASA group (7.06±0.44 cm), and the CIP group (6.85±0.31 cm), as illustrated in Figure 6F.

HE staining of distal colon tissue revealed an absence of lesions in the control group. In contrast, the DSS group exhibited pronounced mucosal structural damage, extensive infiltration of inflammatory cells, and crypt destruction. Treatment with AMPs markedly reduced mucosal ulceration and inflammatory cell infiltration, restored crypt architecture, and clarified tissue structure. The therapeutic efficacy of the 5-ASA group was moderate, whereas the ciprofloxacin (CIP) group demonstrated a comparatively weak therapeutic effect (Figure 6G).

To assess the therapeutic efficacy of each drug on the intestinal inflammatory response, we analyzed the myeloperoxidase (MPO) content in colon tissue, as well as the levels of inflammatory factors and gene expression in both serum and colon tissue. The MPO levels in the DSS group were significantly elevated compared to the control group. In contrast, the MPO levels were markedly reduced in AMPs treatment group, followed by the 5-ASA group, whereas the therapeutic effect observed in the CIP group was comparatively weak (refer to Figure 6H).

### Analysis of inflammatory factors content and gene expression in serum and colonic tissues

Utilizing an ELISA kit (Jianglai Bio), the concentrations of IL-6 and TNF-α were quantified in both serum and colonic tissue samples. The findings indicated that the levels of IL-6 and TNF-α were significantly elevated in the DSS group compared to the control group. Following pharmacological intervention, there was a notable reduction in the inflammatory factor levels across all treatment groups. Specifically, after treatment with AMPs, the inflammatory factor levels in the serum of mice nearly returned to those observed in the control group. Following treatment with 5-ASA and CIP, there was a reduction in the levels of inflammatory factors; however, these levels remained significantly different from those observed in the control group (Figure 7A). Insummary, AMP treatment demonstrated a more pronounced inhibition of DSS-induced proinflammatory cytokine production compared to 5-ASA and CIP. RT-qPCR analysis revealed that the expression levels of IL-6 and TNF-α genes in the colon tissues of the DSS group were significantly elevated relative to the control group. Post-treatment with the drugs, there was a notable decrease in the expression levels of these inflammatory factor genes. In comparison to 5-ASA and CIP, the administration of AMPs resulted in a significant reduction in the expression levels of IL-6 and TNF-α. The efficacy of 5-ASA treatment was observed to be intermediate, while CIP treatment exhibited the least effect. Consequently, AMPs treatment markedly diminished the gene expression of inflammatory markers in colon tissue, as illustrated in Figure 7B. Furthermore, the structural integrity of the colon was restored following pharmacological intervention. The expression levels of tight junction (TJ) proteins in the colon were evaluated using qRT-PCR and Western blot analyses. The findings indicated that the protein levels of ZO-1, Claudin-1, and Occludin were significantly reduced in the DSS group compared to other groups. Subsequent pharmacological intervention led to an upregulation of colonic TJ protein levels in UC mice, with AMPs exhibiting the most pronounced effect, followed by 5-ASA and CIP treatments, as illustrated in Figure 7C.

**Figure 7.**
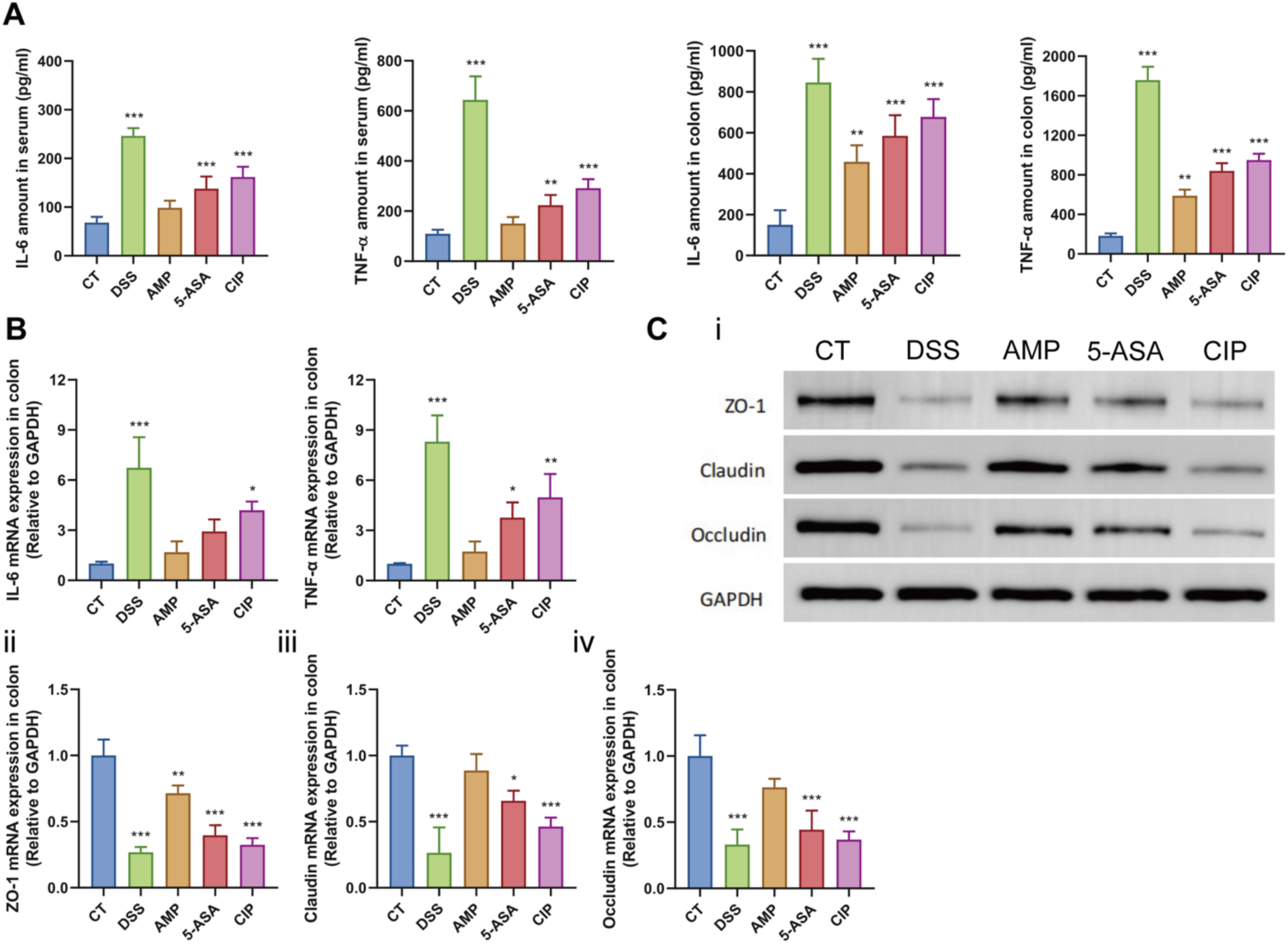
The influence of various therapeutic drugs on clinical symptoms. (A) Inflammatory factors in serum and colon. The data were mean ±SD, n = 6 in each group, and significant differences were indicated by *p < 0.05, **p < 0.01, ***p < 0.001 (analysis of variance). (B) The mRNA expression of inflammatory factors in colon tissues was detected by qRT-PCR. (C) The expression of TJ protein in colon was increased in each group after treatment. Western blot was used to detect the expression of ZO-1, Occludin and Claudin-1 proteins in colon tissue, and qRT-PCR was used to detect mRNA expression. Data are presented as mean ± SD (n = 6); * means p < 0.05, ** means p < 0.01, *** means p < 0.001.

### Analysis of transmission electron microscope results. (AMPs can change the morphology of bacterial cell membrane)

*E. coli* and *S. aureus* treated with PBS, AMPs, and ciprofloxacin were subjected to transmission electron microscopy (TEM) analysis. TEM revealed morphological alterations and cell membrane damage induced by AMPs and CIP. The results indicated that the *E. coli* and *S. aureus* specimens in the control group, as depicted in Figure 8A, exhibited intact and smooth surfaces, characterized by a clear and complete double-layer membrane structure, with a cytoplasmic composition. relatively uniform and dense. Following treatment with AMPs, as illustrated in Figure 8B, the *E. coli* surface transformed into a bubble-like layer, featuring numerous vesicle-like structures that detached from the surface. Additionally, the cytoplasm displayed significant heterogeneity, resembling bacterial vacuolation. Nevertheless, the vesicle-like structure is absent in *S. aureus* (Figure 8C). Following treatment with AMPs, the bacterial cell membrane exhibits significant alterations, appearing rough, wrinkled, and distorted, with the demarcation of the double membrane becoming indistinct. In severe cases, the bacterial cell membrane is compromised, leading to the leakage of intracellular contents. Notably, when both bacterial types were treated with CIP, it was observed that *E. coli* did not exhibit a vesicle-like structural layer (Figure 8D). Upon examining the results of the ultrathin sectionanalysis, it was observed that the primary alterations included the presence of a double-layer membrane structure, indistinct boundaries, leakage of intracellular contents, and complete vacuolization of the bacteria. To elucidate the differential outcomes of antimicrobial peptides (AMPs) on two bacterial types, we hypothesize that these structural changes may underlie the selective affinity of AMPs for distinct bacterial species. This selectivity could also account for the varying bactericidal effects of AMPs on Gram-positive and Gram-negative bacteria.

**Figure 8.**
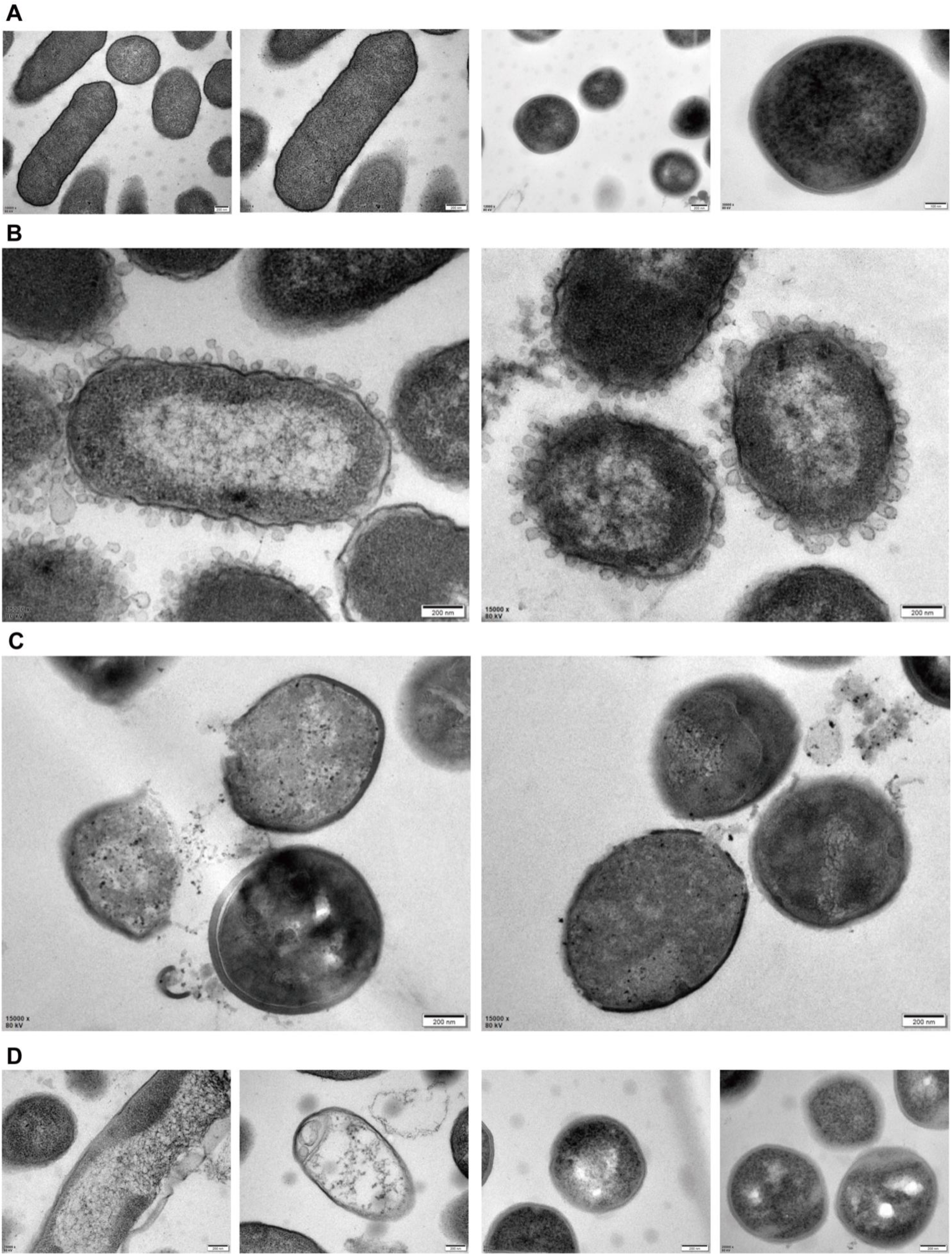
Morphology of bacteria under transmission electron microscope. (A) Under normal circumstances, *E. coli* and *S. aureus* form. (B) Morphology of *E. coli* treated with LR. (C) Morphology of *S. aureus* treated with LR. (D) Morphology of *E. coli* and *S. aureus* treated with CIP.

### The intestinal flora structure and diversity of UC mice were changed after treatment

To determine whether the therapeutic effects of the drug were associated with alterations in the gut microbiota of mice with ulcerative colitis (UC), 16S rRNA sequencing was conducted. The rarefaction curve, rank abundance curve, and species accumulation boxplot indicated that the diversity within the samples was sufficiently captured by the sequencing efforts. Principal Coordinates Analysis (PCoA), utilizing Weighted UniFrac distance, was performed, revealing a significant separation of the DSS group from the CT and AMPs groups. This finding suggests distinct intestinal flora compositions among the groups (Figure 9A). Microbial abundance and diversity assessed through α diversity analyses, including Chao 1, Good’s coverage, and the Shannon index (Figure 9B). Thefindings indicated that the rich diversity of gut microbiota was altered following LR treatment. Furthermore, species distribution histograms were employed to examine the distribution of intestinal flora species. At the phylum level, the dominant taxa were identified as Firmicutes, Bacteroidetes, Verrucomicrobiae, and Proteobacteria (Figure 9C). The treatment with AMPs, 5-ASA, and CIP resulted in an increased relative abundance of Firmicutes and Verrucomicrobia. At the class level, the main class identified were Bacilli, Verrucomicrobiae, Clostridia, and Bacteroidia. At the family level, the predominant families identified were Prevotellaceae, Alistipes, Akkermansiaceae, and Bacteroides. In the DSS group, there was a decrease in relative abundance of Lactobacillus, whereas the relative abundances of Prevotellaceae and Alistipes increased. Furthermore, in the AMPs, 5-ASA treatment groups, the relative abundances of Alistipes, Prevotellaceae and Bacteroidaceae, which are positively correlated with the progression of ulcerative colitis (UC), were reduced (Figure 9C). Notably, within the treatment groups, a marked increase in Akkermansiaceae was observed specifically in the AMPs group. Recent research indicates that the abundance of \**Akkermansia muciniphila*\*, as determined through metagenomic analysis, is inversely correlated with IBD, obesity, and diabetes.^41^ In summary, the treatment group of AMP has the potential to alter the composition of the gut microbiota by decreasing the prevalence of pathogenic bacteria linked to the development of ulcerative colitis (UC) and enhancing the abundance of beneficial bacteria. This intervention ultimately contributes to the partial restoration of the intestinal microbial ecological imbalance induced by DSS.

**Figure 9.**
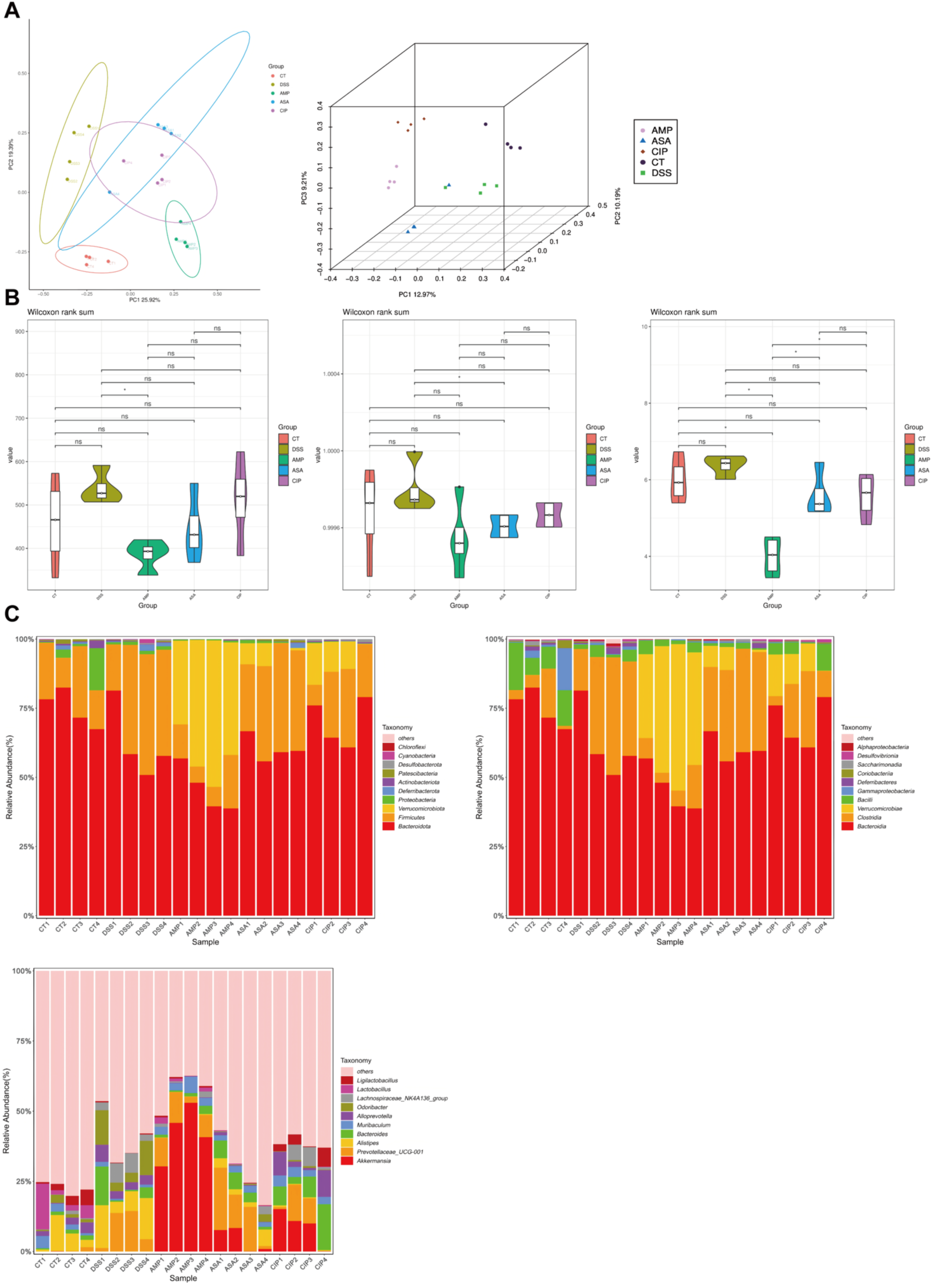
Changes in the structure of microorganisms. (A) principal coordinate analysis (PCoA) graph (B) alpha diversity violin plot (Chao 1, Good’ s coverage and Shannon index). Values are shown as the mean ± SD, n = 4. Significant differences are denoted as *p < 0.05, **p < 0.01, and ***p < 0.001. (C) Classification composition analysis of phylum level class level and family level.

### AMPs can alleviate UC by regulating metabolic pathways

Considering the potential role of intestinal microbiome disturbances in the pathogenesis of UC, we conducted a metabolomic analysis of murine stool samples. Orthogonal Partial Least Squares - Discriminant Analysis (OPLS-DA), a supervised statistical approach for discriminant analysis, was employed. This model facilitates the determination of the Variable Importance in Projection (VIP) scores for both sample sets, thereby elucidating each variable’s contribution to the classification process. The OPLS-DA model demonstrates a robust and reliable performance, evidenced by an R² value exceeding 0.9, indicating significant cluster separation between the AMPs and DSS groups of mice (Figure 10A). The differential abundance of metabolites between these two groups is illustrated by volcano plots (Figure 10B). Analysis of the differential metabolite data reveals alterations in several metabolites between the DSS and AMPs groups, which may be associated with the progression of UC. Furthermore, following treatment with antimicrobial peptides (AMPs), there was a significant upregulation of metabolites such as deoxycholic acid, glycolithocholic acid, and taurodeoxycholic acid. Functional predictions derived from KEGG pathway analysis indicated that AMPs treatment is associated with alterations in purine metabolism, bile acid metabolism, and glycerophospholipid metabolism. These pathways may represent underlying mechanisms contributing to the efficacy of AMPs (Figure 10C). Collectively, these findings suggest the presence of a distinct intestinal metabolome in mice mice with UC, and that administration of AMPs can modulate aberrant intestinal metabolites inthis model.

**Figure 10.**
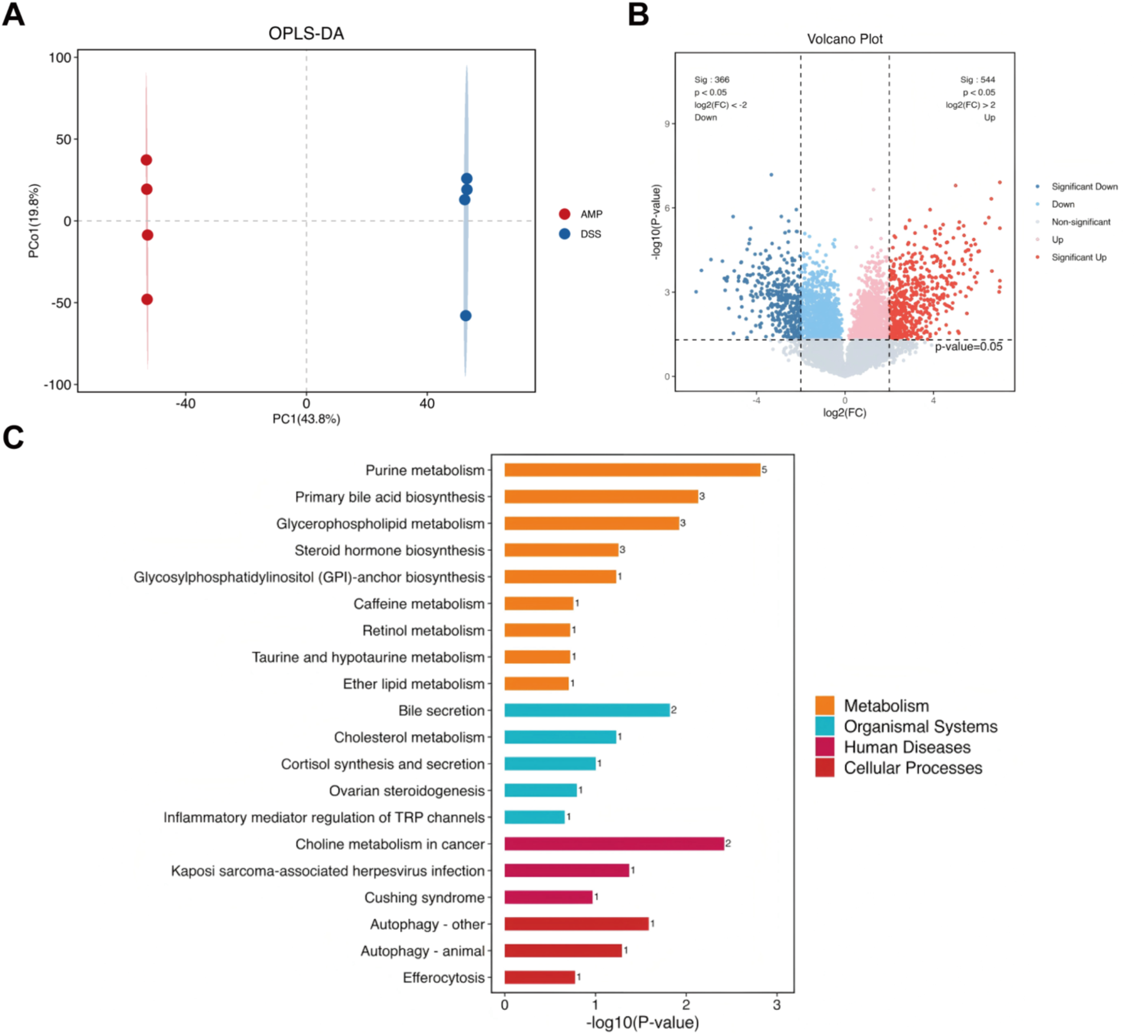
Quantitative analysis of fecal metabolomics was performed. (A) orthogonal partial least squares discriminant analysis (OPLS-DA) plot. Values are Presented as the mean ± SD, n = 4. (B) Volcanic maps showed the accumulation of differences and significant changes in metabolites between the AMPS and DSS groups. (C) Metabolic pathway enrichment analysis of differential metabolites was performed based on KEGG database.

### *Akkermansia muciniphila* replenishment can treat UC

Based on the 16S rRNA sequencing results, we can see that *Akkermansia muciniphila* have a significant advantage in the AMPs treatment group. Therefore, we used *Akkermansia muciniphila* to replenish and treat UC. The results showed that after *Akkermansia muciniphila* replenishment treatment, the body weight of the diseased mice increased, the disease activity index decreased, the spleen index decreased, and the colon length increased. The control group and healthy group mice showed no significant changes in all indicators (Figure 11). In summary, *Akkermansia muciniphila* can alleviate the symptoms of ulcerative colitis mice to some extent.

**Figure 11.**
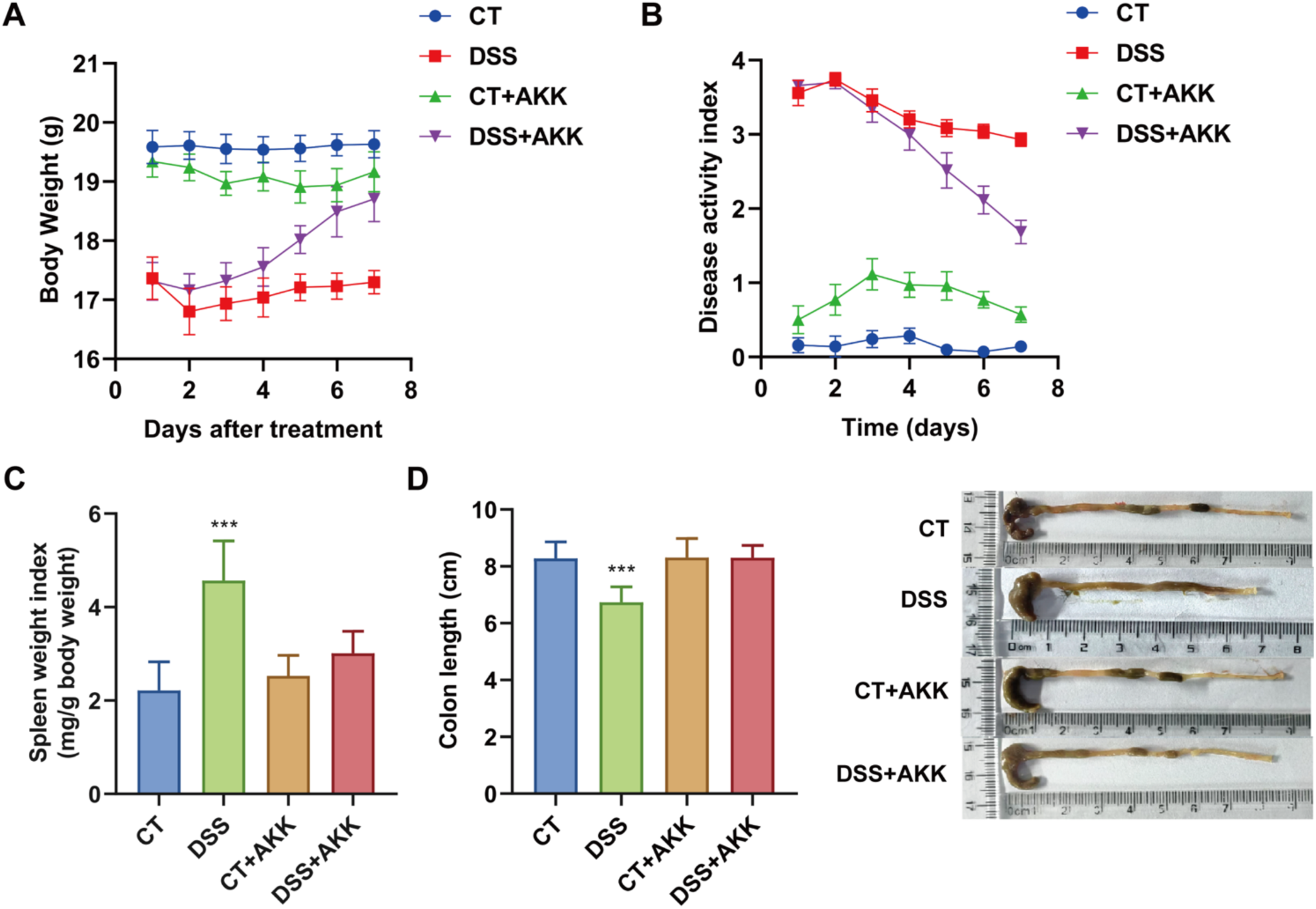
Therapeutic effects of AKK replenishment in UC mice. (A) Changes in body weight of mice after drug treatment. (B) Changes of DAI score in each group of mice after treatment. (C) Changes of spleen index in mice after drug treatment. Significant differences were indicated by *p < 0.05, **p < 0.01, ***p < 0.001 (D) length of colon between the ileocecal junction and the proximal rectum.

## Disscusion

The development of our predictive models, Peptide_predictor and Toxic_predictor, has been a significant endeavor in the field of AMPs research. Our models have demonstrated considerable strengths, with Peptide_predictor showing high accuracy in identifying AMPs, a crucial capability for the rapid screening of potential candidates in large datasets. The optimization process has notably enhanced the F1 score for both models, indicating a balanced improvement in precision and recall, which are pivotal metrics in classification tasks. The comprehensive set of descriptors from the modlamp library has been instrumental in characterizing peptide sequences, contributing significantly to the models’ predictive power. Furthermore, the implementation of 5-fold cross-validation has ensured that our models’ performance is consistent and reliable across different subsets of the data.

However, our models also face certain limitations. The Toxic_predictor, while integral for assessing the safety of potential AMPs, has encountered challenges in accurately predicting toxicity.^42^ This is a critical factor in the practical application of these peptides, as their toxicity profiles can significantly influence their therapeutic utility. Additionally, the observed correlation between toxicity and bactericidal activity has complicated the model’s ability to distinguish between toxic and non-toxic peptides, potentially due to overlapping features that are indicative of both properties.

To address these challenges and further enhance our models, we plan to explore more advanced machine learning architectures, such as machine learning models, which could potentially capture more complex patterns in the data and improve toxicity prediction.^43–44^ We also intend to refine our feature set, possibly by incorporating domain-specific knowledge or alternative descriptor sets that could better differentiate between toxic and non-toxic peptides. Implementing strategies to address class imbalance, such as oversampling or adjusting class weights, could also improve the model’s ability to predict less frequent outcomes, like toxicity.

Furthermore, we aim to integrate additional data sources, such as peptide structure information or experimental toxicity data, which could provide the models with a more comprehensive understanding of peptide properties. Establishing a framework for continuous model evaluation as new data becomes available will ensure that the models remain up-to-date and relevant. By addressing the current limitations and exploring new avenues for model development, we aim to enhance the predictive capabilities and applicability of our models in the field of antimicrobial research, ultimately contributing to the discovery and development of novel AMPs with optimized safety profiles.

In the current study, we demonstrate that algorithmically screened AMPs can reduce DSS induced intestinal inflammation, and that this beneficial effect may come from improving the inflammatory state and restoring the loss of the intestinal epithelial barrier. In addition, our study showed that after drug treatment in each group, the therapeutic effect of AMPs was relatively significant, highlighting the effectiveness of algorithmically screened AMPs in the treatment of UC. NF-κB pathway is a very important pathway in cell signaling, and it plays a key role in regulating various physiological processes such as immune response and inflammatory response. This pathway regulates the expression of multiple genes in response to a variety of stimuli, including pro-inflammatory cytokines, free radicals, bacterial and viral infections. Studies have shown that activation of a component of the NF-κB pathway produces pro-inflammatory cytokines, such as TNF-α, in UC mice, leading to inflammation of the colon.^45–46^ Consistent with this finding, we found elevated levels of inflammatory factors in serum and colon tissue in the DSS group, and decreased levels of TNF-α and IL-6 after treatment, while in our in vitro cellular anti-inflammatory experiments, we found that amp reduced LPS-induced cellular inflammation and reduced levels of TNF-α and IL-6.

At present, studies have shown that intestinal mucosal barrier dysfunction also plays an important role in the pathogenesis of UC.^47^ It was found that the physical structure of intestinal mucosal epithelial tight junction (TJ) changed and intestinal mucosal barrier dysfunction appeared in UC patients,^48–49^ which was consistent with our experimental results, and the expression of tight junction proteins ZO-1, Claudin-1 and Occludin significantly decreased in DSS group. This study found that AMPs increased the expression of tight junction proteins, suggesting that AMPs has a positive effect on restoring the integrity of the colon epithelial barrier.

We also found that AMPs can regulate the composition of gut microbiota, with a significant increase in the proportion of *Akkermansia muciniphila* (AKK) in the stool of mice in the AMPs treated group. In addition, our AKK replantation experiment results also proved that AKK replantation could alleviate the symptoms of ulcerative colitis. AKK, which is mainly colonized in the intestine, is known to improve intestinal immune function and is important for maintaining normal intestinal function.^50–51^ Studies have found that AKK can improve a variety of intestinal diseases,^52–53^ and the abundance of AKK was significantly reduced in mice with intestinal inflammatory diseases such as IBS, UC,^54^ appendicitis^55^ and allergic diarrhea.^56^ Moreover, studies have also shown that AKK is associated with weight gain, decreased expression of pro-inflammatory cytokines, and restoration of intestinal epithelial barrier function in DSS induced colitis.^57–58^ Therefore, Based on the results of our experiments, we hypothesized that AMPs can restore colon barrier damage by regulating intestinal flora composition and increasing AKK.

Changes in small molecule metabolites affected by AMPs are also an important part of UC therapy. The results of stool metabolomics in mice in this study showed significant changes in the levels of deoxycholic acid, glycolithocholic acid, taurodeoxycholic acid and other metabolites. Our research results show that the algorithm model we established has good prediction results, and the selected LR can induce intestinal flora structure rearrangement, regulate bile acid metabolism, and repair intestinal mucosal barrier, which provides a theoretical basis for the development of novel AMPs therapy for UC.

### Limitations of the study

While our findings highlight the therapeutic promise of machine learning-driven antimicrobial peptides (AMPs) in ulcerative colitis (UC), several limitations merit consideration. First, the inherent "black-box" nature of machine learning models limits mechanistic interpretation of how specific physicochemical features govern AMP functionality, despite their predictive accuracy. Second, the translational relevance of our DSS-induced murine colitis model remains constrained by inherent differences between murine and human gut microbiome composition and immune responses. Third, while 16S rRNA sequencing provided insights into microbial shifts, metagenomic or metatranscriptomic analyses would better resolve functional alterations in low-abundance taxa and strain-level dynamics. Fourth, our study focused on short-term therapeutic efficacy; long-term safety assessments of AMPs, including potential off-target effects on commensal microbiota or epithelial integrity, remain to be addressed. Finally, although *Akkermansia muciniphila* enrichment correlated with AMP efficacy, causality has been partially validated through *Akkermansia muciniphila* replenishment treatment, the underlying mechanisms remain to be fully explored. These limitations underscore the need for expanded preclinical models, multi-omics integration, and rigorous safety profiling to advance AMPs toward clinical translation.

## Materials and methods

Phosphate-buffered saline (PBS, powder), agar powder((C12H18O9)n), sodium dodecyl sulfate (SDS,C12H25SO4Na), triton X-100 (C34H62O11), were purchased from Beijing Solarbio Science & Technology Co., Ltd. (Beijing, China); yeast extract and tryptone were purchased from Thermo Fisher Scientific Co., Ltd. (USA); nutrient broth (NB) and brain heart infusion (BHI) broth were purchased from Qingdao Hope Biotechnology Co., Ltd. (Qingdao, China); 2,2,2-Trifluoroethanol (TFE) was purchased from Sigma-Aldrich. (USA); sodium chloride (NaCl), absolute ethanol, 25% aqueous glutaraldehyde solution were purchased from Sinopharm Chemical Reagent Co., Ltd. (Shanghai, China); dimethyl sulfoxide (DMSO) was purchased from J&K Chemical Technology (China); Cell Counting Kit-8 (CCK8) was purchased from GlpBio (USA); Trizol, HiScript III RT SuperMix for qPCR and Taq Pro Universal SYBR qPCR Master Mix were purchased from Vazyme (China), Mouse TNF-α ELISA kit and Mouse IL-6 ELISA kit were purchased from Jiang lai Bio (China), ZO-1 Rabbit pAb, Claudin 1 Rabbit mAb, Occludin Rabbit pAb and GAPDH Antibody were obtained from ABclonal (China); Calcein/PI Cell Viability/Cytotoxicity Assay Kit was purchased from Beyotime (China). 5-Aminosalicylic Acid, Artificial gastric fluid (sterile), Artificial intestinal fluid (sterile) were purchased from Shanghai yuanye Bio-Technology Co., Ltd. (Shanghai, China); Ciprofloxacin (CIP) was purchased from Sigma-Aldrich (USA). Human red blood cells (hRBCs) and serum were sourced from individuals in good health.

All mice utilized in this study were procured from Jinan Pengyue Experimental Animal Breeding Co., Ltd. (Jinan, China) and maintained under specific pathogen-freeconditions.

### Dataset trained machine learning model for screening AMPs

#### Construction of data sets

##### Antimicrobial Peptide Dataset Construction

To develop a comprehensive dataset of antimicrobial peptides (AMPs), we commenced by collecting data from publicly accessible AMP repositories, specifically the Antimicrobial Peptide Database (APD) and the Database of Antimicrobial Peptides from Spiders (dbAMPs). Following the amalgamation of these datasets, we eliminated redundant entries and excluded peptides that exhibited solely anticancer activity, as well as those exceeding 60 amino acids (AA) in length. This process yielded a final set of 13,637 unique AMP sequences, with lengths ranging from 5 to 50 AA.

##### Non-antimicrobial Peptides Dataset Construction

Concurrently, we curated a dataset of non-antimicrobial peptides (non-AMPs) from the UniProt database. To ensure the integrity of the dataset, we applied the ‘Subcellular Location’ filter to select entries specific to the cytoplasm while excluding those associated with antimicrobial, antibiotic, antiviral, antifungal, effector, or excretory substances.^29^ Following the elimination of duplicate sequences and those identical to entries in the AMPs dataset, we retained a total of 9,385 non-AMPs sequences ranging in length from 5 to 50 amino acids..

#### Optimization of model construction and screening of AMPs

##### Descriptor Description

Subsequently, we utilized the modlamp library’s modlamp. descriptors module to compute a variety of descriptors for the amino acid sequences.^30^ We employed all descriptor scales available within this module and computed autocorrelation descriptors for each sequence, configuring the calculate_autocorr parameter to a value of 5. These descriptors facilitated a quantitative assessment of the sequences’ physicochemical properties and structural attributes.^31^

##### Model Construction

Subsequently, we utilized the ‘RandomForestClassifier’ from the ‘sklearn’ library to develop a random forest classifier for sequence classification tasks.^32–34^ During the model training phase, critical hyperparameters were configured to enhance performance. Specifically, the ‘max depth’ parameter was set to 10 to limit the maximum depth of the trees, ‘max_features’ was configured to ‘auto’ to enable automatic feature seletion, and ‘n_estimators’ was fixed at 100 to specify the number of trees in the forest. All other hyperparameters were maintained at their default settings.

Model performance was assessed using metrics such as accuracy, precision, recall, and the F1 score. To ensure the obustness of the evaluation, a 5-fold cross-validation procedure was employed.

##### Hyperparameter Optimization

We systematically fine-tuned the hyperparameters of the random forest classifier to enhance model performance. A comprehensive hyperparameter search space was delineated, encompassing the maximum depth of the trees (max_depth) with values ranging from 5 to 20, feature selection strategies (max_features) including auto selection, square root, and log2 methods, as well as the number of trees in the forest (n_estimators) varying from 10 to 100. Furthermore, we evaluated the minimum number of samples necessary to split an internal node (min_samples_split) within the range of 1 to 10, as well as the allocation of class weights (class_weight) for imbalanced datasets, opting for balanced weights and balanced subsampling strategies.

Employing scikit-optimize (skopt), we systematically explored hyperparameter combinations within this predefined search space to identify the optimal configuration that maximizes model accuracy, precision, recall, and F1 score. This process was accompanied by a 5-fold cross-validation to ensure robustness and generalizability of the results. ^35–36^

##### Antimicrobial Peptides Screening

We utilized a genetic algorithm in conjunction with a trained prediction model to screen potential AMPs.^37^ The initial population comprised 6,000 sequences, with an equal distribution of known AMPs and non-AMPs to ensure diversity and variability.

During each iteration of the genetic algorithm, the prediction model was employed to preliminarily evaluate the sequences, assessing their antimicrobial and toxicity properties. Subsequently, the charge and hydrophobicity of the sequences computed utilizing the modlamp.descriptors module. The screening criteria targeted sequences exhibiting a charge range of 5 to 7 and hydrophobicity between 40% and 60%, attributes linked to the efficacy of AMPs.

The selected sequences were then subjected to crossover operations, emulating natural selection and genetic recombination processes. To preserve genetic diversity and circumvent local optima, 3000 novel sequences were introduced. Following ten rounds of iteration, 22 target sequences that met the specified criteria were selected from a pool of 6,373 candidate sequences, thereby enhancing the screening efficiency and increasing the likelihood of discovering novel AMPs. Next, we used the Antimicrobial Peptide Database (APD) to search for natural AMPs targeting intestinal pathogens, and modified them. After the modification, we used the algorithm for screening.

### In vitro characterization of antimicrobial peptides

#### Synthesis and Structural Parameter Characterization of AMPs

##### Synthesis of AMPs

The aforementioned AMPs were synthesized by GL Biochem (Shanghai, China) utilizing the standard solid-phase FMOC method, with amidation at the C-terminus. All synthesized AMPs underwent purification via high-performance liquid chromatography, achieving a purity exceeding 95%. The molecular weights of the AMPs were verified through mass spectrometry (MS). Subsequently, all AMP samples were stored at −80 °C.

##### Prediction of AMPs’ Structural Parameters

The molecular weights of all AMPs were determined using the Novopro tool (https://novopro.cn/tools/).The net charge and hydrophobic content of the peptides were calculated utilizing the prediction tool available in the APD (https://aps.unmc.edu/AP/). Secondary structure predictions of the AMPs were conducted using AlphaFold3 (https://alphafoldserver.com/).

#### 2.3.2. Bactericidal validation of antimicrobial peptides screened by algorithm in vitro

##### Bacterial cultivation

In this study, three types of pathogens were selected: the typical Gram-negative bacterium *Escherichia coli* (*E. coli*), the typical Gram-positive bacterium Staphylococcus aureus (*S. aureus*) and the multidrug-resistant bacterium Acinetobacter baumannii (*AB-29*). The bacterial cultures were maintained in Luria-Bertani (LB) medium. Individual bacterial colonies were incubated at 37 °C with agitation at 180 rpm for 12 hours, resching the logarithmic growth phase for subsequent use.

##### In vitro antimicrobial activity assay

The minimal bactericidal concentration (MBC), defined as the lowest concentration of an antimicrobial agent required to kill bacteria, was determined via the colony-count assay.^38–39^ The bacterial suspension was diluted with phosphate-buffered saline (PBS) to a concentration of 10^5^ cfu/mL. Antimicrobial peptides (AMPs) were prepared in PBS at concentrations of 400 μg/mL, 200 μg/mL, 100 μg/mL and 50 μg/mL. The bacterial suspension was then mixed in a 1:1 ratio with the AMPs and co-incubated for 2 hours,before being inoculated onto solid medium for further analysis. The Minimum Bactericidal Concentration (MBC) was assessed by quantifying the number of colonies formed on a solid medium. A survival rate histogram was generated using GraphPad Prism 8.

#### In vivo biocompatibility determination of antimicrobial peptides screened by algorithm

##### Cell cultivation

RAW 264.7 macrophages were cultured in Dulbecco’s Modified Eagle Medium (DMEM) supplemented with 10% fetal bovine serum (FBS) and seeded in 6-well plates at a density of 1.0 × 10^5^ cells per well. Similarly, mouse colon cancer cells (CT-26.WT) were cultured in RPMI 1640 medium with L-Glutamine also supplemented with 10% FBS, and seeded inoculated in 6-well plates at a density of 1.0 × 10^5^ cells per well.

##### Cell viability

Murine macrophages (RAW264.7) and mouse colon cancer cells (CT-26) were cultured to the logarithmic growth phase and subsequently seeded into 96-well plates at a density of 10^5^ cells per well. The cells were incubated in a cell incubator maintained at 37 °C with 5% CO2 for 24 hours. Following this incubation period, the supernatant was aspirated, and varying concentrations of AMPs were prepared in DMED and 1640 media, achieving final concentrations of 12.5, 25, 50, 100, and 200 μg/mL. These preparations were then added to the respective wells, and the cells were incubated for an additional 12 hours. Wells without AMPs served as blank controls. Subsequently, 10 μL of CCK-8 solution was added to each well after removing the supernatant, followed by further incubation for another 2 hours. Finally, the absorbance was measured at 450 nm using a microplate reader (Thermo Fisher Scientific, Multiskan MK3, China). The cell survival rate was calculated using GraphPad Prism 8.

##### Hemolytic activity assays

Fresh blood was collected and centrifuged, the supernatant was discarded, and the erythrocytes were washed with PBS 4-6 times until the supernatant was clear. A 2% v/v erythrocyte suspension in PBS was then mixed with AMPs at final concentrations of 12.5, 25, 50, 100, and 200 μg/mL. Samples treated with PBS served as the negative control, while those treated with 2% Triton X-100 served as the positive control. Following a 2-hour incubation period at 37°C under gentle conditions, the supernatant was collected via centrifugation.,The absorbance of the supernatant at 570 nm was subsequently measured using a microplate reader (Thermo Fisher Scientific, Multiskan MK3, China).

A histogram depicting the hemolysis rate was generated using GraphPad Prism 8.

##### Cell life and death staining experiment

Murine macrophages (RAW264.7) and mouse colon cancer cells (CT-26) were cultured to the logarithmic growth phase and then seeded into 6-well plates, which were placed in a cell incubator maintained at 37°C with 5% CO2 for 24 hours. The supernatant was removed, and different concentrations of AMPs were added to DMEM and 1640 media, respectively, to achieve final AMP concentrations of 25, 50, 100, and 200 μg/mL. The cultures were incubated for 12 hours. Samples without AMPs served as blank controls, while cells treated with 75% alcohol for 30 minutess served as positive controls. Following the removal of the supernatant, the cells were gently washed three times with PBS. An appropriate volume of Calcein AM/PI (Beyotime) test solution was then added, and the cells were incubated at 37℃ for 30 minutes in the dark. Post-incubation, the staining was observed using a fluorescence microscope (Nikon Eclipse Ti2, Japan).

#### Anti-inflammatory assays

Initially, a macrophage inflammation model was established using RAW264.7 macrophages stimulated with lipopolysaccharide (LPS) at concentrations of 0.1 μg/ml, 1 μg/ml, 10 μg/ml and 100 μg/ml. Cell viability was assessed using the CCK-8 assay. Subsequently, samples without AMPs and LPS served as negative controls, while samples with only LPS served as positive controls, Additionally, 0.1 μg/ml LPS was incubated with AMPs at concentrations of 25 μg/ml, 50 μg/ml, 100 μg/ml, and 200 μg/ml for 12 hours. The supernatant was then collected, and the levels of inflammatory cytokines were quantified using IL-6 and TNF-α assay kits.

#### Circular Dichroism Spectroscopy Detection of AMPs

The secondary structures of the AMPs were investigated using a circular dichroism (CD) spectrometer (Jasco J-1500, Tokyo, Japan). To mimic various environmental conditions, PBS (100 mM, pH7.4) was employed to replicate a normal physiological environment, 50% trifluoroethanol (TFE) was used to simulate a hydrophobic environment, and a 30mM sodium dodecyl sulfate (SDS) solution was utilized to represent a microbial membrane environment. The AMPs were dissolved in each of these three solvents to a final volume of 1 mL and a concentration of 200 μg/mL. The CD spectra were recorded in a 1 mm path-length quartz cell using the aforementioned CD spectrometer. The scanning procedure was conducted three times, and peak plots were generated using GraphPad Prism 8.

#### Stability of antimicrobial peptides in vitro

##### In vitro thermal stability test

Both AMPs solution and the blank control PBS solution were incubated in a water bath at 60℃ for 2 hours. The AMPs were subsequently diluted with PBS to achieve final concentrations of 60 μg/ml, 80 μg/ml and 100 μg/ml. The final concentration of *E. coli* and *S. aureus* was standardized to 10^5^ CFU/mL. The AMPs solution was then mixed in a 1:1 ratio with the bacterial suspensions of *E. coli* and *S. aureus*, respectively. Following this, the mixtures were inoculated onto solid media, to determine the minimum bactericidal concentration (MBC). A survival line graph was constructed using GraphPad Prism 8.

##### In vitro acid-base stability assays

PBS solutions with pH values of 2 and 8.4 were prepared using HCl (1 mol/L) and NaOH (1 mol/L), respectively. AMPs were then dissolved in these PBS solutions with varying pH values and incubated at 37℃ for 2 hours. Following the incubation period, the pH of the AMPs was adjusted to 7.4. The subsequent procedure adhered to the sterilization protocol utilized in the thermal stability test. The minimum bactericidal concentration (MBC) was calculated, and the survival curve was generated using GraphPad Prism 8.

##### In vitro proteolytic stability assays

The AMPs were dissolved in gastric juice and small intestine fluid. Pepsin and trypsin were diluted with PBS to achieve a final concentration of 1 mg/ml. The mixture was then incubated at a constant temperature of 37℃ for 2 hours. Following the incubation period, the subsequent procedure adhered to the sterilization protocol outlined in the thermal stability test. The minimum bactericidal concentration (MBC) was calculated, and a survival curve was generated using GraphPad Prism 8.

### In vivo characterization of antimicrobial peptides

#### Animal experiments

To investigate the effect of AMPs on ulcerative colitis (UC), an animal experiment was conducted in accordance with international, national, and institutional guidelines. All applicable institutional and governmental regulations concerning the ethical use of animals were strictly adhered to. Healthy female C57BL/6J mice, aged 6−8 weeks and weighing 18−20 grams, were randomly assigned to five groups, each consisting of six mice. The groups included a control group, a DSS-induced colitis group, an AMPs treated group (5 mg/kg), a 5-ASA treated group (5 mg/kg), and a CIP treated group (5 mg/kg). The mice were acclimatized for one week prior to their inclusion in the study. The control group received PBS throughout the entire duration of the experiment. The experimental groups were administered 3% DSS for 7 days. Daily measurements included mouse body weight, fecal collection from each group, and the performance of an occult blood test. From day 8 onward, the DSS group received PBS, while the AMP group, 5-ASA group, and CIP group were treated with AMP, 5-ASA, and CIP, respectively. Seven days post-treatment, the mice were weighed, anesthetized, and blood was collected from the canthal inner canthi. Additionally, the colon and spleen were harvested for further analysis.

#### Mice treatment-related indicators

##### Disease activity index evaluation (DAI)

The disease activity index (DAI) of ulcerative colitis (UC) was evaluated According to the grading system specified in Table S1^59^. Daily scores were computed for each mouse by averaging body weight, stool consistency, and the extent of intestinal bleeding.

**Table S1.** Disease activity Index score sheet^59^.

| Score | Weight loss | Stool consistency | Blood |
| --- | --- | --- | --- |
| 0 | none | normal | negative hemocult |
| 1 | 1-5 % | soft but still formed | negative hemocult |
| 2 | 6-10 % | soft | positive hemocult |
| 3 | 11-18 % | very soft; wet | blood traces in stool visible |
| 4 | >18 % | watery diarrhea | gross rectal bleeding |
| 5 | death |  |  |

##### Histopathological analysis

For histological assessment of UC, colon samples were fixed in 4% paraformaldehyde and subsequently embedded in paraffin. Paraffin-embedded sections were dewaxed to water and immersed in a high-resolution constant dye pretreatment solution for one minute.Subsequently, the tissue sections were immersed in hematoxylin dye solution for a duration of 3-5 minutes, gollowed by differentiation with a differentiation solution, and subsequently treated with a blueing solution to revert the sections to a bluehue. Post-dehydration in 95%ethanol, eosin staining was conducted for 15 seconds. The sections were then subjected to further dehydration using a series of alcohols and xylene, and finally sealed with neutral gum. Microscopic images were acquired using a Nikon Eclipse Ti2 microscope (Nikon, Japan).

##### MPO activity measurement

MPO (Myeloperoxidase, MPO) activity was assessed in homogenized mouse colon tissues prepared in PBS buffer. Following homogenization, the samples were centrifuged, and the supernatant was collected for analysis. MPO activity in the colon was assessed ultilizing commercially available kits. The analysis was conducted at a wavelength of 460 nm using a visible spectrophotometer.

##### Determination of inflammatory factors in serum and colonic tissue

The concentrations of inflammatory cytokines, interleukin-6 (IL-6) and tumor necrosis factor-alpha (TNF-α), in both serum and colon tissue were quantified using specific assay kits. Absorbance measurements were recorded at 450 nm with a microplate reader (Thermo Fisher Scientific, Multiskan MK3, China). The cytokine concentrations were determined from a standard curve and subsequently plotted using GraphPad Prism 8 software.

##### RT-qPCR assays

Total RNA was extracted from colonic tissues utilizing Trizol (Genstone Biotech) according to the manufacturer’s protocol. First-strand cDNA synthesis was conducted using reverse transcription kits (Q712-02, Vazyme). Real-time PCR was performed employing SYBR Green Master Mix reagents (#22208, Tolobio) and primer mixtures (Table S2). The mRNA expression levels of target genes were normalized to GAPDH and quantified using the ΔΔCt method.

**Table S2.**
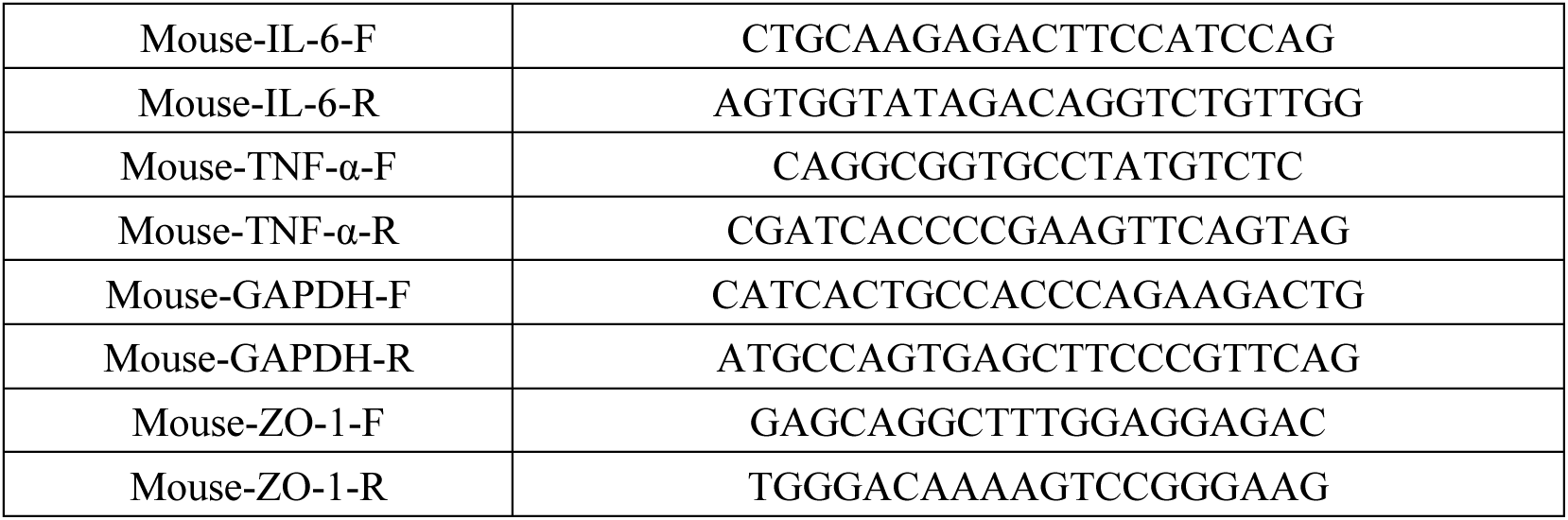
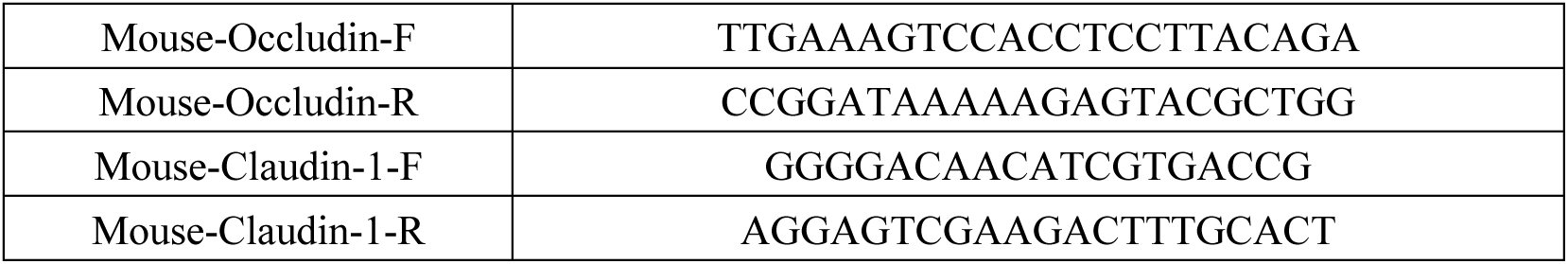
qRT-PCR primers.

##### Western blot analysis

Colonic tissues were lysed using RIPA buffer supplemented with protease inhibitors (Solarbio, China). Protein concentrations were quantified employing a BCA kit (Solarbio, China), and subsequently normalized across all experimental groups. To optimize protein separation, gels composed of polyacrylamide and sodium dodecyl sulfate (SDS) were prepared with varying concentrations, tailored to the molecular weights of the target proteins. SDS-PAGE was performed, followed by the electro transfer of proteins onto polyvinylidene fluoride (PVDF) membranes. The membranes were then blocked with 5% skim milk at room temperature for 2 hours. Shortly thereafter, the samples were incubated with the primary antibody overnight at 4℃. The PVDF membrane was subsequently washed with PBST, and incubated with the secondary antibody for 1 hour. A color-developing solution was then applied to the PVDF membrane, and images were captured. Bands were visualized and quantified using GAPDH as a reference.

### Transmission electron microscope analysis

#### Sample pretreatment

*E. coli* and *S. aureus* suspensions treated with AMP, antibiotics, and PBS were subsequently washed and fixed with 2.5% glutaraldehyde.

#### transmission electron microscope

The pre-treated samples were initially washed with PBS buffer. Subsequently, the samples were dehydrated using a graded series of acetone and ethanol. Following dehydration, the samples were immersed in a penetrant solution consisting of 50% acetone and 50% embedding agent for one hour. The samples were then transferred into EPON812 epoxy resin embedding agent and incubated at 37℃, 45℃ and 60℃ for 24 hours to facilitate heating polymerization, Post-polymerization, the samples were sectioned using an ultra-thin microtome. The sections were stained with lead nitrate and uranium acetate. Morphological changes were examined using a transmission electron microscope (TEMJSM JEOL-840).

### 16S rRNA gene sequencing

#### DNA extraction and amplification

Fecal samples from the experimental mice were collected and stored at −80 °C prior to analysis. Total genomic DNA was extracted utilizing a Fecal Genomic DNA Extraction Kit (TianGen), adhering to the manufacturer’s protocol. The concentration and integrity of the DNA were assessed using a NanoDrop 2000 spectrophotometer (Thermo Fisher Scientific, USA) and agarose gel electrophoresis. The extracted DNA was subsequently stored at −20°C until further processing. This DNA served as the template for PCR amplification of bacterial 16S rRNA genes, employing barcoded primers and Takara Ex Taq polymerase (Takara). The 16S rRNA genes were amplified for diversity analysis utilizing primers 343F and 798R.

#### Library construction and sequencing

The quality of the amplicons was assessed via agarose gel electrophoresis. Subsequently, the PCR products were purified using AMPure XP beads (Agencourt) and subjected to an additional round of PCR amplification. The final amplicons were quantified employing the Qubit dsDNA Assay Kit (Thermo Fisher Scientific,USA). The concentrations were then normalized for sequencing. Sequencing was conducted on an Illumina NovaSeq 6000 platform, generating 250 bp paired-end reads (Illumina Inc., San Diego, CA; OE Biotech Company, Shanghai, China).

### Metabolomic analysis

#### Sample pretreatment

For non-targeted metabolite analysis, fecal samples stored at −80°C are first thawed at room temperature. The thawed fecal samples are then homogenized using a methanol-water mixture. Following pre-cooling, the samples are subjected to grinding at 60 Hz for 2 minutes. Ultrasonic extraction is performed in an ice water bath. Subsequently, the samples are centrifuged, and the supernatant is extracted using a syringe, transferred to an LC sample vial, and stored at −80℃. Quality control (QC) samples are prepared by pooling aliquots from all individual samples.

#### LC-MS/MS analysis

Metabolomic data analysis was conducted by Shanghai Luming Biological Technology Co., LTD (Shanghai, China). Metabolomic profiling was performed utilizing an ACQUITY UPLC I-Class Plus system coupled with a Thermo QE mass spectrometer. Data acquisition was carried out in both positive and negative ion modes.

#### Data Preprocessing and Analysis

The LC-MS data were processed using Progenesis QI V2.3 software (Nonlinear Dynamics, Newcastle, UK) for baseline filtering, peak identification, integration, retention time correction, peak alignment, and normalization. Differential metabolites were subsequently analyzed for KEGG pathway enrichment (http://www.genome.jp/kegg/).

### *Akkermansia muciniphila* replenishment experiment

Using the method mentioned earlier to construct the UC model, the mice were divided into four groups: the healthy group (CT), the control group (CT+AKK), the disease group (DSS), and the AKK treatment group (DSS+AKK). The healthy group and the disease group were treated with PBS by gavage, while the control group and the AKK treatment group were treated with AKK bacteria by gavage. The treatment lasted for 7 days, after which various indicators of the mice were detected.

## Resource availability

### Lead contact

Further information and reasonable requests for resources and reagents should be directed to and will be fulfilled by the lead contact, Pengfei Cui.

### Materials availability

Requests for materials should be made via the lead contact. All unique/stable reagents generated in this study are available from the lead contact without restriction.

## Data and code availability

- ***Data***: All data reported in this paper will be shared by the lead contact [Pengfei Cui] upon reasonable requests.
- ***Code***: All original code will be shared by the lead contact [Pengfei Cui] upon reasonable requests.
- ***Additional Information***: Any additional information required to reanalyze the data reported in this article is available from the lead contact upon request.

## Acknowledgments

This work was supported by National Natural Science Foundation of China (No. 31902421).

## Author contributions

Conceptualization, P.C., H.M. and Z.W.; Methodology, H.M., Z.W., P.C., H.M., S.C., J.W., H.Y., Y.L. and Z.G.; Formal Analysis, H.M., Z.W., Z.G., H.M., J.W. and P.C.; Investigation, H.M., Z.W., P.C., H.M., S.C., J.W. and Z.G.; Software, Z.W.; Resources, P.C. Writing – Original Draft, H.M., H.M. and P.C.; Writing – Review & Editing, H.M., H.M. and P.C.; Funding Acquisition, P.C.; Supervision, P.C.

## Declaration of interests

The authors declare no competing interests.

## Notes

### Competing Interest Statement

The authors have declared no competing interest.

